# Landscape of oncoviral genotype and co-infection via Human papilloma and Hepatitis B viral tumor in-situ profiling

**DOI:** 10.1101/2020.03.24.006130

**Authors:** Adrian Bubie, Fabien Zoulim, Barbara Testoni, Brett Miles, Marshall Posner, Augusto Villanueva, Bojan Losic

## Abstract

Hepatitis B virus (HBV) and human papillomavirus (HPV) infection are known risk factors for developing several cancers. However, the effect of viral genotype and co-infection in actually driving oncogenesis remains unclear. We have developed and deployed a new scalable, high throughput tool (ViralMine) for sensitive and precise oncoviral genotype deconvolution using tumor RNA sequencing data from 537 virally infected liver, cervical, and head and neck tumors. We provide the first comprehensive integrative landscape of tumor-viral gene expression, viral antigen immunogenicity, patient survival, and mutational profiling organized by tumor onco-viral genotype. We find that HBV and HPV genotype, and surprisingly high rates of multi-genotype co-infection, serve as significant predictors of patient survival, tumor immune responsiveness, and *APOBEC* activity modulation. Finally, we demonstrate that HPV genotype strongly associates with viral onco-gene expression over cancer type, implying expression may be similar across episomal and stochastic integration-based infections.

**Significance Statement:** We demonstrate that scalable, high-accuracy oncoviral genotyping, gene expression, and co-infection estimation is feasible from legacy tumor RNA-seq data. While HBV and HPV genotype are known risk factors for oncogenesis, viral genotype and co-infection are shown to strongly associate with disease progression, patient survival, mutational signatures, and putative tumor neoantigen immunogenicity, facilitating novel clinical associations with infections.

## Introduction

Chronic infection with Hepatitis B virus (HBV) and Human Papillomavirus (HPV) are well known oncogenic risk factors, with strong viral genotype associations (*1, 2*). Given that HBV related hepatocellular carcinoma (HCC) and HPV related head and neck and cervical cancer incidence is on the rise globally (*3*–*5*), scalably exploiting tumor RNA sequencing (RNA-seq) data to accurately infer detailed viral signatures is clinically urgent. While averaged infection phenotypes such as viral load and predominant genotype have been previously characterized and shown to be strong prognostic factors in cancer development (*6*–*10*), the effects of more granular measures such as exon-level viral expression or the ratio of expressed viral genotypes (co-infection) have not yet been fully mapped out in the host tumor microenvironment. This leaves key facets of these DNA oncoviral infections unknown, creating a clinical blindspot for the development of potential new anti-oncoviral therapeutic options.

Here we present a new *in situ* tool to comprehensively characterize DNA oncoviruses, which we applied to 1230 tumor samples spanning across liver, cervical, and head and neck cancers. Although HBV and HPV infect highly disparate cancers via different mechanisms, their strong genotype specific association with oncogenic risk, relatively unknown co-infection rates (*11*–*13*), and disease progression associations can be naturally combined in an integrative, viral genotype-centric, study leveraging tumor RNA-seq data. Indeed, our tool ViralMine, extracts and quantifies viral RNA from high-depth coverage tumor sequencing with high fidelity sequence recovery, allowing for precise and accurate viral genotype deconvolution and viral exon-level expression analysis (**Figure 1A**) within the context of the tumor microenvironment. While previous studies have adopted similar negative selection strategies to extract viral sequences from tumor RNA profiling (*10, 14*–*16*), our data-driven method goes considerably further by facilitating scalable viral gene expression characterization and deconvolution of complex viral co-infection patterns.

**Fig. 1:**
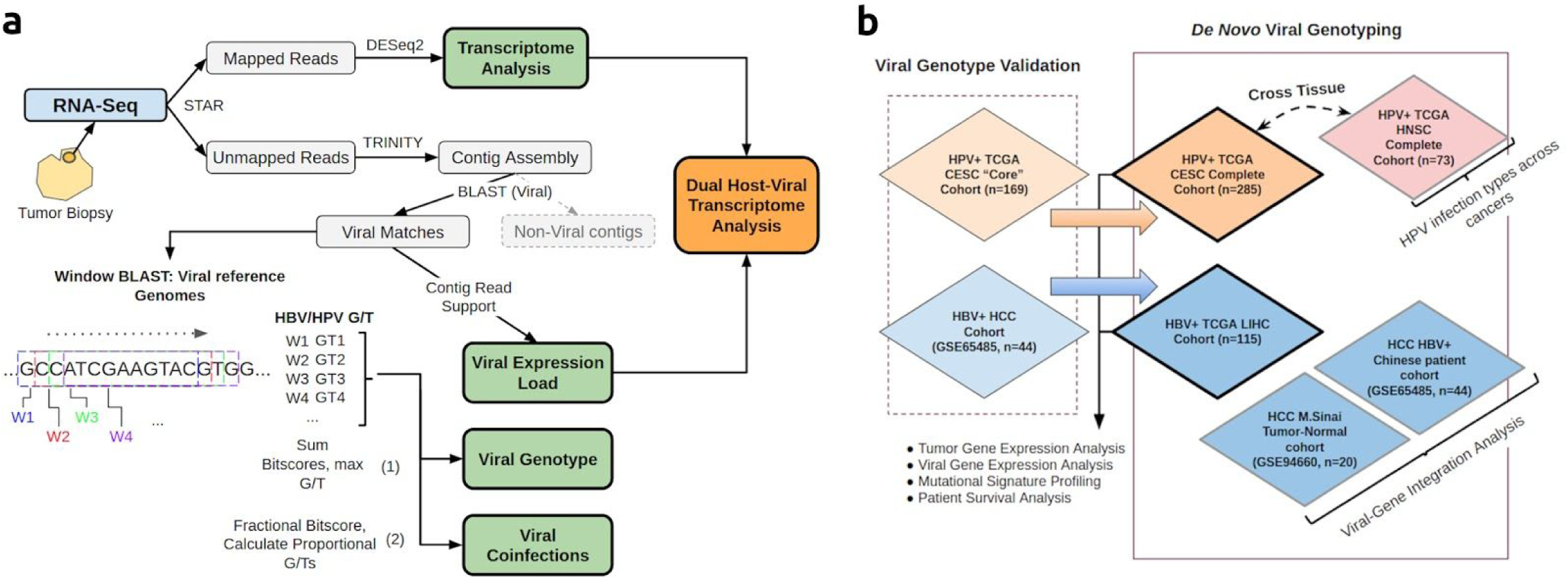
ViralMine workflow and study design. (**A**) Workflow summary of our tool, ViralMine. RNA sequencing reads unaligned to the human reference are assembled into contigs and aligned against viral reference databases. Virus genotype, tumor co-infection status and viral expression are quantified using alignment and contig read support information. (**B**) Validation and discovery cohorts for viral genotyping. Sample groups included in each analyses indicated.

We found that viral genotype is strongly associated with overall patient survival, cancer specific expression signaling, and tumor immunogenicity in HCC. Similarly, viral genotype was linked to significant expression differences in host oncogenesis and immune response pathways in HPV related cervical cancer, and striated patient survival. We found that HPV co-infection creates a notable increase in average putative neoantigen immunogenicity, indicating a potential emergent property of co-infection within cervical tumors. Finally, a comparison of HPV gene expression in head and neck cancers and cervical cancers revealed significant variation across viral tumor genotype but not cancer type.

## Results

### ViralMine: in-silico cross-cohort genotyping validation and co-infection discovery

In order to validate the key genotyping capability of ViralMine (**Figure 1A**, see methods), we obtained tumor RNA-seq data for the “core set” of cervical cancers from the TCGA with previous HPV infection information (n=178, 169 HPV+) and a group of 50 Chinese HCC patients screened for HBV infection (GSE65485, n=50, 44 HBV+) (**Figure 1B**) (*15, 17*), for a total of 213 virally infected samples. We inferred viral genotypes in both cohorts against the existing genotype information, obtaining perfect sensitivity in both datasets, and specificities of 0.97 and 0.95 in the HPV+ cervical cancers and HBV associated HCC tumors, respectively **(Supplemental Figure 1**). These results demonstrate that the overall performance of ViralMine in the cervical TCGA (CESC) core set cohort nearly perfectly matched consensus calls made using a combination of different assays, including MassArray, BioBloom Tools and PathSeq RNA-Sequencing inference techniques (*15*). We also observed robust average performance using read downsampling with patient randomization, indicating even relatively low viral expression is sufficient for accurate genotyping with ViralMine (**Supplemental Figure 9**), though we do note that immunotherapy treatments which stimulate viral clearance may have an adverse effect on viral sequence recovery (**Supplemental Figure 10**).

Applying ViralMine to large-scale tumor expression datasets, especially those with missing or incomplete viral information, we exhaustively screened liver cancer patients from the TCGA (LIHC, n=334) for HBV infection and found 115 positive patients. This included 44 patients previously reported HBV+ (*18*), with the majority (85/115, 74%) infected with HBV genotype C (**Figure 2A,B**). We also screened a different dataset of 21 HCC patients for combined HBV integration and genotyping analysis (**Figure 1B; Figure 2B**) (GSE94660, n=21), and found 11 patients positive for HBV genotype C, 9 HBV for genotype B, and 1 patient HBV- (*19*). Similarly, we analyzed HPV infected tumors across the entire set of cervical cancers from the TCGA (CESC, n=304), finding 285 HPV+ (93.7%), with HPV genotypes for the virally infected tumors indicated in **Figure 1C**. Among the cervical histological subtypes, neither HPV+ adenocarcinomas nor squamous cell carcinomas were found to be correlated with a specific viral genotype (**Supplemental Figure 2A**). We also screened cancers from the head and neck TCGA cohort (HNSC, n=521), finding 73 HPV+ (14%), of which the vast majority (81%) were HPV16, which restricted inter-tumoral genotype comparisons (**Figure 2A,B**).

**Fig. 2:**
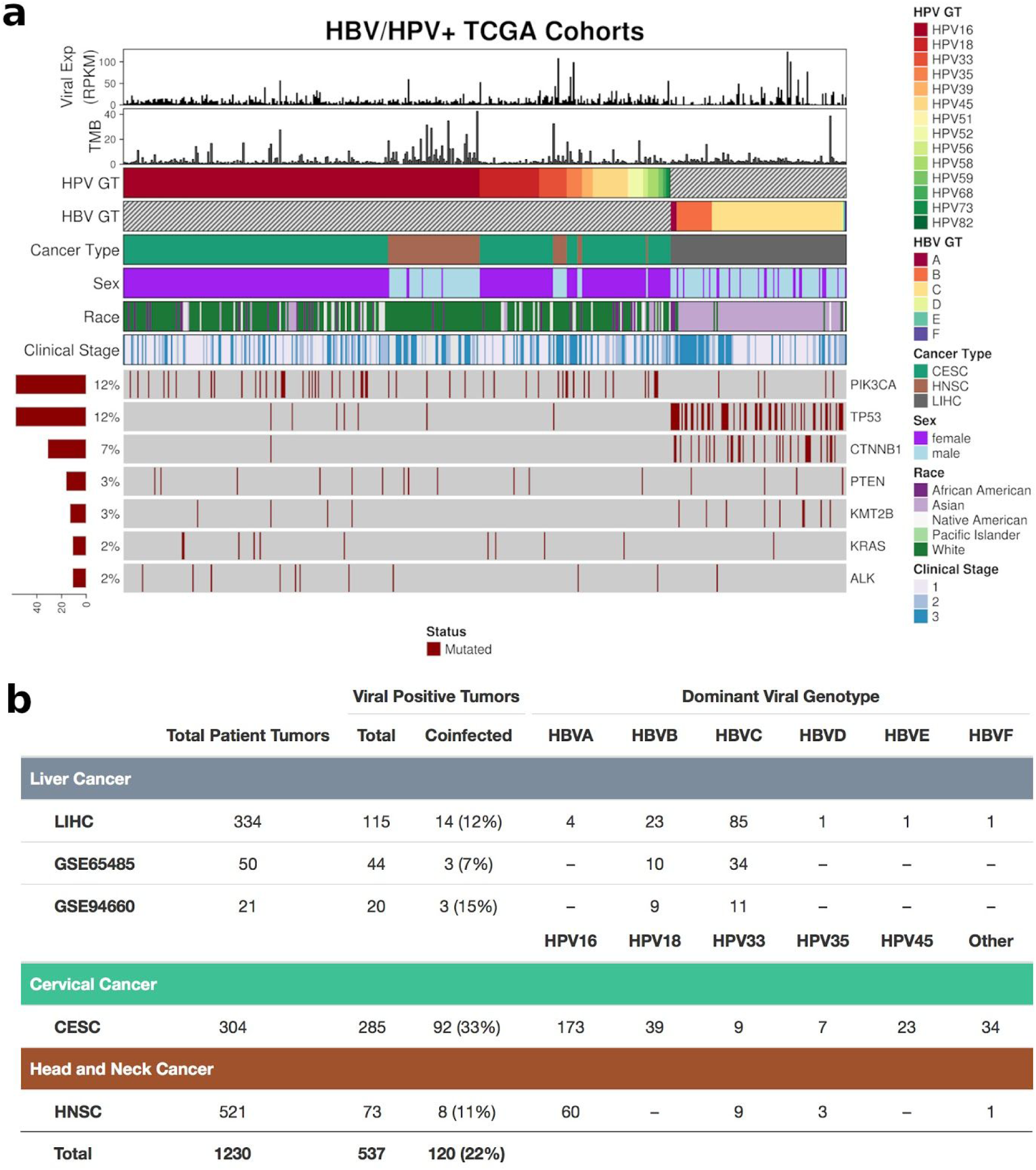
Cohort and Oncoviral infection overview. (**A**) Viral genotype, expression, and key clinical phenotype and somatic mutation information for the three TCGA HPV and HBV cohorts. Percentage patients with denoted somatic mutation indicated by barplot. (**B**) Summary table of viral infections by genotype across cohorts.

Co-infections, defined by expression of more than one genotype supported by at least 10% of the total viral sequence alignment, of HPV genotypes were found in 92 of the 285 HPV+ cervical cancers (32.5%) with 82 infected with two HPV genotypes (29%), and 10 with three HPV genotypes (3.5%), yielding higher co-infection rates compared to previous co-infection surveys, considering much smaller cohorts of cervical lesions (**Figure 2B, Supplemental Figure 3A**) (*11, 12*). As with primary HPV infection genotypes, these surprisingly high co-infection rates among CESC samples were not preferentially associated with either adenocarcinomas or squamous cell carcinoma subtypes (**Supplemental Figure 2B**). In the TCGA head and neck HPV associated tumors, 8 of 73 were co-infected (11%) with two HPV genotypes (**Figure 2B; Supplemental Figure 3B**). In HBV-related HCCs in the TCGA cohort, 14 tumors were co-infected with more than one HBV genotype (12%), of which 12 had two genotypes and 2 had three concurrent genotypes (**Figure 2B; Supplemental Figure 3C**). While it is technically possible to conflate a recombinant virus with distinct viral genotypes only looking at RNA data, typical recovered viral contigs (average of 500-1000bp) were of a length to suggest that we are quantifying the latter.

We also noted a lack of correlation between patient viral load and somatic tumor mutational burden (TMB), and between viral genotype and viral load, across cancer type (**Figure 2A; Supplemental Figure 4**). This indicated viral genotype was not conflated with other molecular covariates and represents an independent tumor read-out. However, we found no significant associations between viral genotype and several expression based immune activity markers for either HPV or HBV, potentially signalling viral genotype alone does not have a significant effect on tumor immune activity.

### HBV genotype C associated with molecular signatures of aggressive HCC

In order to test the hypothesis that particular viral genotypes are associated with unique downstream onco-expression signaling, we performed gene set enrichment analysis (GSEA) of the differentially expressed genes between HBV genotype C (n=77) and HBV genotype B (n=18) associated HCCs of TCGA. We found HBV genotype C tumors were enriched in pathways involved in cell proliferation, tumor recurrence, and tumor growth, while cell survival pathways and liver specific genes sets were downregulated versus patients with HBV B tumors (FDR<0.0001) (**Figure 3A**; **Supplemental Table 1**). Interestingly, for tumors with HBV co-infection (n=14), genes in pathways involved in 3’ UTR translation, peptide chain elongation, and miRNA deregulation were significantly downregulated compared to tumors with a single HBV genotype (n=100; FDR<0.0001)(**Supplemental Figure 5**; **Supplemental Table 1**), indicating regulatory instability may increase with multi-genotypic HBV infections in the tumor. Integration analysis using only expressed transcripts among HBV+ genotype C and B HCC patient groups (n=172) identified HBV C preferred integration loci in known HCC driver genes *CCNE1* and *KMT2B* (*18, 20*), while total average rate of integration among the two tumor groups (1.81 integrations per HBV C tumor, 1.85 per HBV B tumor) was similar (two-tailed t-test, p=0.9)(**Supplemental Figure 6**). Finally, while the *APOBEC3* pathway has been shown to activate in response to HBV infection in liver malignancies (*21*), we did not find any significant differential expression associated with HBV genotype within HBV+ tumors.

**Fig. 3:**
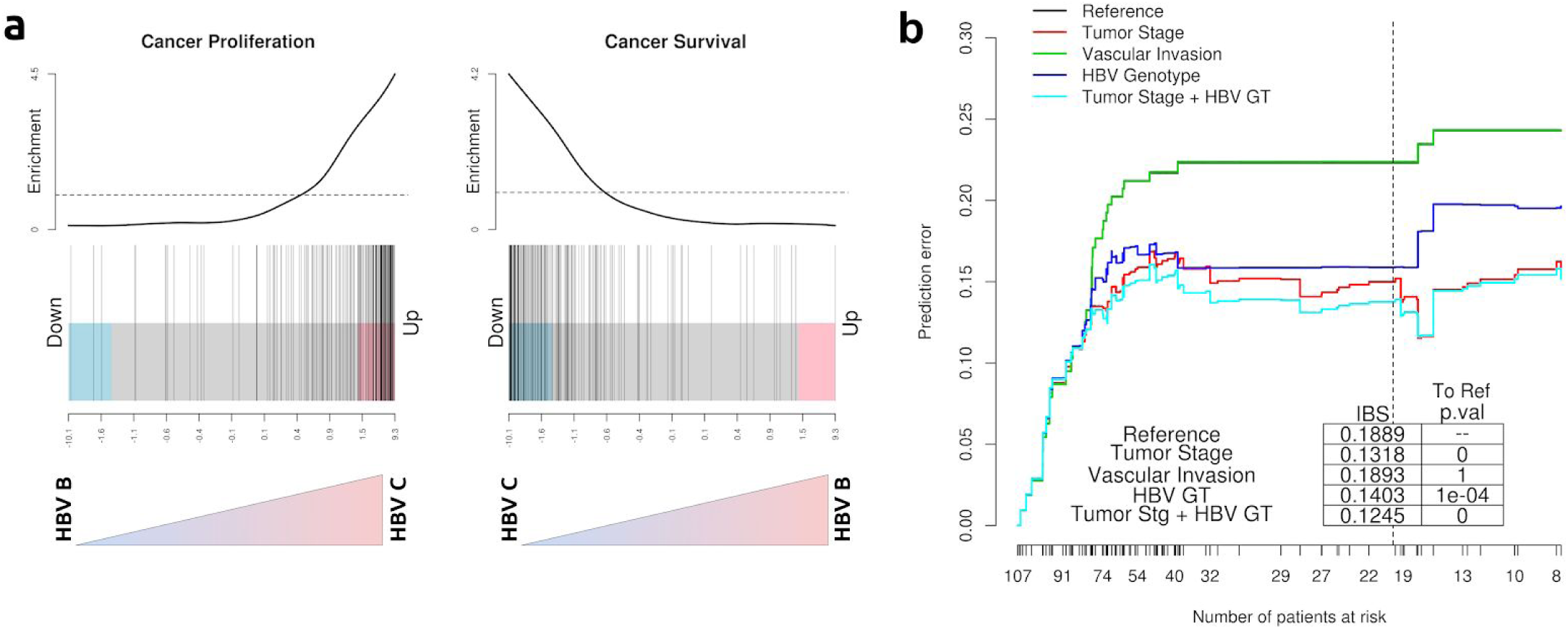
HBV genotype affect HCC expression and outcome. (**A**) Barcode plots indicating gene up- and downregulation in cancer proliferation and survival pathways given HBV genotype. Cancer progression is significantly upregulated and cancer survival significantly downregulated in HBV C-like patients compared to HBV B patients. Bars along the bottom of the plot indicate individual pathway gene enrichment scores. (**B**) Prediction error curves for cox proportional hazard models of LIHC patients from the TCGA. Survival prediction error for each model, listed in the upper left corner, is calculated across the number of patients at any given survival time, and plotted compared to a reference model using no clinical covariates for prediction. Integrated Brier scores (IBS) for each model calculated for survival times encompassing 80% of patient events are listed in the inset table. Significance values comparing each model against the reference curve are indicated next to each score.

Given the genotype driven molecular differences found above, we sought to determine if the predictive impact of viral genotype in patient survival was significant. We constructed Cox proportional hazard models in HBV-related HCC from TCGA and, using bootstrap resampling to control overfitting, computed robust time dependent Brier scores for models using clinical tumor stage, tumor vascular invasion, and tumor HBV genotype. Comparing the resulting prediction error curves (**Figure 3B**), we found both tumor stage and HBV genotype significantly reduced prediction error against the naive reference model, while vascular invasion did not. Survival prediction was additionally improved using a model including both tumor stage and HBV genotype terms, significantly so over the HBV genotype only model (likelihood ratio test, p<0.001) and almost so over tumor stage alone (p=0.101). HBV genotype is a remarkably strong predictor of overall patient survival.

### Cervical cancer HPV genotype and co-infection status differentiate survival and possess unique molecular fingerprint

We surveyed differentially expressed host genes between HPV genotype 16 (HPV16, n=173) and HPV genotype 18 (HPV18, n=39) infected cervical tumors of TCGA, representing the two most dominant genotypes. Via GSEA, we found pathways in tumor vasculature and endothelial growth are enriched in HPV16 over HPV18 infected tumors, while TNF-signal regulated apoptosis is downregulated (FDR<0.01)(**Figure 4A**; **Supplementary Table 1**). Similarly, we tested for the effect of co-infection and found that in contrast to single HPV genotype cervical tumors (n=193), those infected with multiple HPV genotypes (n=92) are enriched in non-*IFR3* antiviral activation of lymphocytes and B-cell antigen activation pathways (FDR<0.05)(**Figure 4B**), suggesting preferential activation of alternative immune regulatory pathways in HPV co-infected tumors. We also carried out viral integration analysis finding no significant sites of preferential integration across either HPV genotype (*16*) or co-infection status. (**Supplemental Figure 7**) (*14, 15, 22*). Finally, while it was previously shown that *APOBEC3* expression (associated with antiviral activity) is upregulated in HPV-associated head and neck cancers but not cervical cancers (*10*), we found that *APOBEC3* is significantly upregulated in cervical tumors with HPV16 over HPV18, and in tumors with a singular HPV infection over those with multiple HPV genotypes, controlling for HPV genotype (HPV18, HPV45) (**Figure 4D**). Thus the increase in *APOBEC* activity seen in HPV+ patients (*10*) is actually further dependent on HPV genotype and co-infection rather than viral infection status alone. Increased *APOBEC3* activation has been shown to be linked with further tumor mutagenesis (*23*), signaling that HPV16 infection may further drive cervical tumorigenesis.

**Fig. 4:**
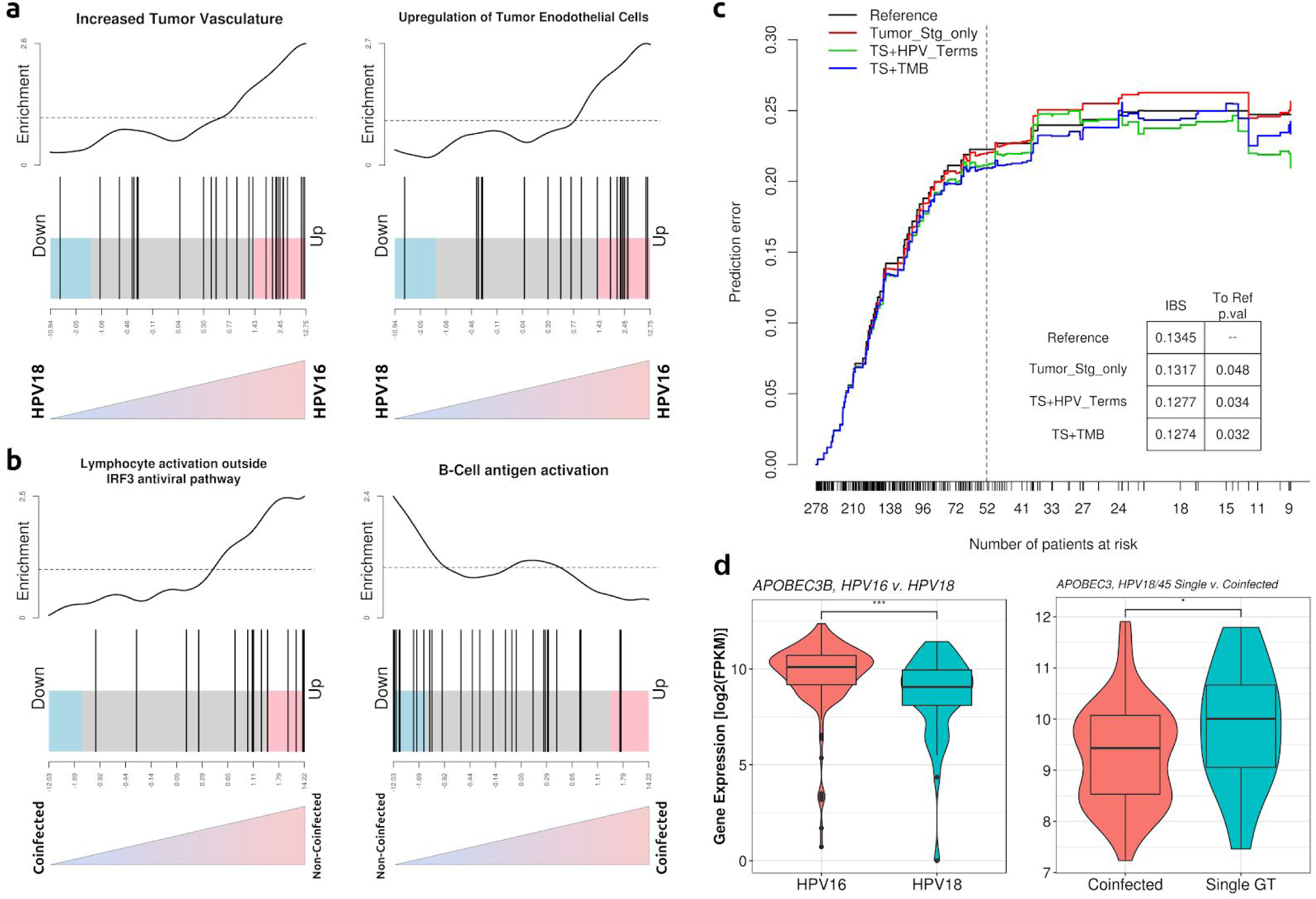
Cervical cancer expression and outcome varies with HPV co-infection and genotype. (**A**) Barcode plots indicating pathway enrichment of tumor vasculature and tumor endothelial cell growth in patients with HPV16 cervical tumors compared with HPV18 infected tumors. Bars indicate individual pathway gene up- and downregulation. (**B**) Cervical cancers with HPV co-infections are upregulated in non-*IRF3* antiviral lymphocyte activation, and downregulated in B-cell antigen activation, compared with tumors with a single HPV infection genotype. (**C**) Prediction error curves for cox proportional hazard survival models of HPV+ cervical cancer patients from the TCGA (CESC), comparing the effect of HPV genotype, viral expression, and co-infection status (*HPV_Terms*) with tumor mutational burden (*TMB*) as tumor clinical stage (*Tumor_Stg_only, TS*) is maintained as a covariate. Brier scores for each model at survival times encompassing 80% of patient events are included in the inset table, with significance values comparing each model with the reference curve. (**D**) Normalized *APOBEC3* expression between cervical tumors with HPV16 and HPV18 (left) infections, and cervical tumors with HPV18 or HPV45 single genotypes against HPV18/45 co-infections (right). Comparison by two-sided Wilcoxon rank-sum test (p_genotype_= 1.4×10^−4^ ; p_co-infection_= 0.05).

In order to quantify the predictive effect of viral genotype and co-infection status on patient survival, we built Cox proportional hazard models of overall survival in the CESC TCGA using predictors clinical tumor stage, HPV molecular terms (tumor HPV genotype, viral expression, and viral co-infection status), and patient TMB. As before, we computed and compared time dependent Brier scores between models to compare prediction error (**Figure 4C**). Although all models slightly but significantly reduced prediction error with respect to the naive reference model, we note that it is driven by the relatively few number of events (deaths) among cervical patients. However, the addition of HPV molecular terms to clinical tumor stage (*TS+HPV_Terms*) significantly reduced prediction error compared to tumor stage alone (*Tumor_Stg_only*) indicating additional predictive survival power is encoded by HPV phenotype (likelihood ratio test, p=0.023). Furthermore, we found this genotype driven model performs on par with a model using tumor stage and patient TMB (*TS+TMB*) (likelihood ratio test, p=1) as predictors. Taken together, our results confirm that HPV16 modestly but significantly associates with poorer survival (*24*) and that HPV phenotype is a reasonable predictor of survival, adding significant predictive power beyond tumor staging and tumor mutation burden alone.

### Viral genotypes associate with distinct mutational signatures and phenotypes

To further parse the association of overall tumor mutation burden and viral genotypes by specific mutation type, we derived single base-pair substitution (SBS) based mutational signatures (*25*) for the LIHC, CESC, and HNSC TCGA by stratifying patients based on tumor HBV genotype and HPV viral clade, respectively. As shown by *Alexandrov et al.*, these signatures are key descriptors of cancer phenotypes and can serve as prognostic and predictive biomarkers (*26*). Using their algorithm (see methods), we found signatures linked to tobacco smoking and those of unknown etiology (see <u>Single Base Substitution (SBS) Signatures</u>) preferentially enriched in the HBV B associated liver cancer mutational profile and absent from the HBV C profile (**Figure 5A;** average enrichments in **Supplemental Table 2**). Both patient groups were enriched in SBS22 and SBS24, linked to carcinogenic aristolochic acid and aflatoxin B1 exposures, respectively, as previously reported enriched across HBV+ LIHC patients (*18*). In cervical cancers, however, we found that tumor mutational profiles associated with HPV a9 infections (including genotypes 16, 31, 33, 35, 52, and 58) are enriched in signatures SBS3, SBS9, SBS26, and SBS29, while these signatures are absent from the HPV a7 (including genotypes 18, 39, 45, 59, and 68) mutational profile (**Figure 5A; Supplemental Table 2**). Signatures SBS3 and SBS26 are both linked to defective DNA damage repair pathways, while SBS29 is associated with exposure to chewing tobacco and SBS9 serves as a signature of hypermutation in lymphoid cells. On the other hand, signature SBS40 (of unknown etiology) is enriched only in HPV a7 tumors. While no head and neck tumors were associated with HPV genotypes in the a7 clade, we found head and neck tumors associated with HPV a9 infections were actually enriched in the majority of the same signatures as the cervical HPV a9 tumors (SBS3, 9, 29 and 36; **Figure 5A; Supplemental Table 2**), with exceptions for signatures SBS36, linked to defective base excision repair, and SBS26.

**Fig. 5:**
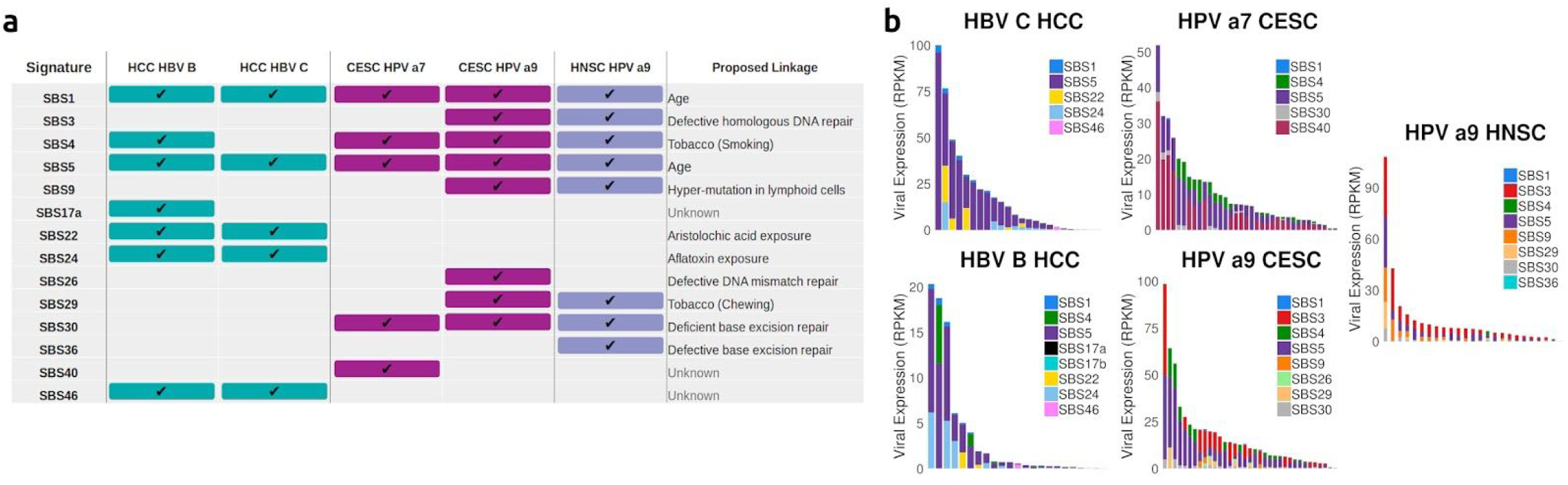
Mutation profile signatures unique to viral genotypes. (**A**) Table of single basepair substitution (SBS) COSMIC global mutational signatures present in mutational profiles of patients in viral genotype subgroups. Colored boxes indicate presence of the SBS signature within the *de novo* mutational profiles for viral genotype groups in the labeled columns. Etiologies for each signature are listed to the left. (**B**) Signature contributions across representative patient mutational profiles, organized by viral expression in the tumor. Each bar represents one patient of the given viral genotype group, with color fill corresponding to the proportion of the patient’s total mutations attributed to the indicated SBS signature.

In order to clearly visualize any trends in mutational signatures scaling with overall viral burden, we selected representative patients spanning the range of total viral expression from each tumor genotype (**Figure 5B**). We found no association between viral expression and mutational signature enrichment in patients, across any HBV or HPV cohorts (p_min_ > ∼0.7), nor across patient TMB (p_min_ > ∼0.5), (**Supplemental Figure 8**). Our results suggest that viral genotype does indeed delineate specific mutational signatures, regardless of total viral expression or tumor mutation burden, across both HPV associated cervical and head and neck cancers and HBV associated HCC liver cancer. Apart from enriching in mutational signatures with well-established functional associations (SBS1, 4, 5, 26), other strong genotype-specific enrichments are for mutational signatures of unknown etiology (SBS17, 40, 46), suggesting potentially new functional viral associations and hypotheses.

### HBV genotype and HPV co-infection drive differential tumor immunogenicity

Given the evidence that HBV genotype serves as a significant predictor of patient survival in HCC, we hypothesized that a lower tumor antigen immunogenicity might drive worse outcomes in HBV C associated HCC patients. Inferring HLA types calculated from HBV+ patient tumor RNA-Seq in the TCGA LIHC, we estimated the MHC-I binding affinities for a total of 37,222 and 142,539 unique tumor neoantigens across HBV B patients and HBV C patients, respectively. Comparing the cumulative distribution functions of the predicted neoantigen MHC-I binding affinities reveals a significantly higher binding affinity bias (lower ic50) for HBV B related tumor neoantigens over peptides from HBV C related HCCs (p<0.0001, **Figure 6A**). Indeed, we found that the ratio of immunogenic tumor mutations (somatic mutations with at least one predicted neoantigen with ic50<500nM) to total tumor mutations is greater among HBV B infected tumors than HBV C tumors (p<0.001; **Figure 6A, inset**), suggesting that there is both a higher occurrence of immunogenic mutations in HBV B HCCs, and, consequently, TIL recruitment likelihood by tumor neoantigens from these tumors is greater on average than in HBV C related tumors.

**Fig. 6:**
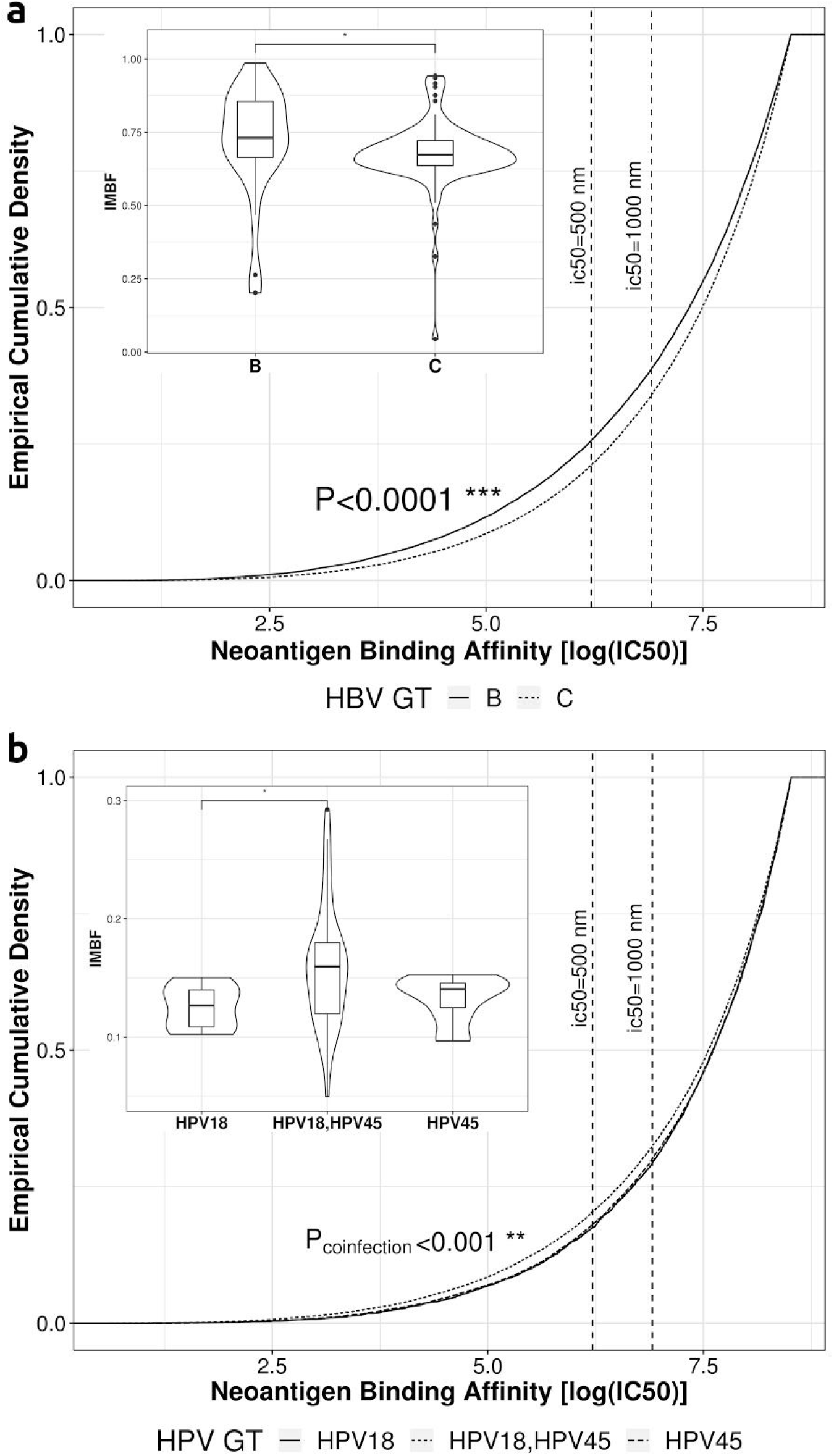
Viral genotype and co-infection modulate tumor neoantigen immunogenicity. (**A**) Tumor neoantigen MHC-I binding affinity cumulative density functions for LIHC TCGA patients infected with HBV B (solid line) and HBV C (dashed line). Smaller values of ic50 indicate stronger binding affinity of tumor neoantigen with T-cell MHC-I complex. Dotted lines labeled at ic50 of 1000 and 500 demarcate “weak” and “strong” antigen binding affinity thresholds, respectively. Significance by one-sided KS-test indicates cumulative density function of HBV B neoantigens is significantly greater (more immunogenic) than HBV C neoantigens. Inset box-and-violin plot compares the immunogenic mutation burden frequency (IMBF), the ratio of mutations generating at least one strongly immunogenic neoantigen (ic50<500) to total mutational burden, between HBV B associated tumors and HBV C tumors. (**B**) Neoantigen MHC-I binding affinity cumulative density functions for CESC TCGA patients with cervical HPV18 (solid line) and HPV45 (long dash line) infection and HPV18/45 (dotted line) co-infection. Ic50 binding affinity thresholds marked as above. Significance by one-sided KS-test indicates co-infected tumor neoantigen cumulative density function is significantly greater than either neoantigen binding affinity distribution for tumors with single HPV infections. Inset box-and-violin plot compares IMBF between the three non- and co-infected groups.

In the TCGA CESC cohort, we tested whether HPV co-infection was associated with a modulated effect on tumor antigen immunogenicity. Controlling for HPV genotype, we compared the predicted tumor neoantigen binding affinities of cervical cancers infected with HPV18, HPV45, and those with co-infections of both HPV18 and HPV45 (totals of 5,108; 13,289; and 37,995 antigens, respectively). Tumor neoantigens from tumors with HPV co-infections had greater average binding affinity over tumors with either single HPV type alone (HPV18 vs HPV18/45, p<0.001; HPV45 vs HPV18/45, p<0.001; **Figure 6B**), while there was no significant difference in binding affinity distributions for singly infected tumors (HPV18 vs HPV45, p=0.8). The ratio of immunogenic mutations to total TMB was found to be higher in co-infected tumors than in both HPV18 and HPV45 tumors (**Figure 6B, inset**), though only significantly compared to HPV18 (p=0.038; p=0.12), while there was no significant difference in ratios between HPV18 and HPV45 patients (p=0.45). In summary, we find intriguing preliminary in-silico evidence suggesting a differential immunogenicity among tumors infected across HBV genotypes and HPV co-infection.

### HPV gene expression is more closely correlated with genotype than to cancer type

In order to examine the effect of cancer cell-type in viral expression from the genotype-specific perspective, we compared cervical and head and neck cancer at both the total HPV expression level and across HPV genes by leveraging high-quality viral whole-transcript recovery from ViralMine (**Figure 7**). While there was no statistically significant difference in total HPV expression between HPV clades across cervical and head and neck tumors (**Figure 7, panel 9**), we found normalized HPV gene expression for cervical tumors with HPV infections in the a7 (n=73) clade is lower across *E1, E2, E5, E7*, and *L1* genes compared to tumors with HPV a9 (n=205) infections (**Figure 7, panels 1, 2, 4, 6, 7;** p_max_ < 0.05). Crucially, HPV gene expression is significantly upregulated in head and neck tumors with a9 clade infections (n=69) over CESC a7 tumors in the *E1, E2, E5*, and *E7* HPV genes (**Figure 7, panels 1, 2, 4, 6;** p_max_ < 0.05), indicating HPV genotype may actually have a stronger effect on the expression in these specific viral genes than cancer cell-type. It is important to note that we could not include HNSC tumors with HPV a7 infections in our comparisons as no TCGA HNSC were infected with HPV genotypes in the a7 clade.

**Fig. 7:**
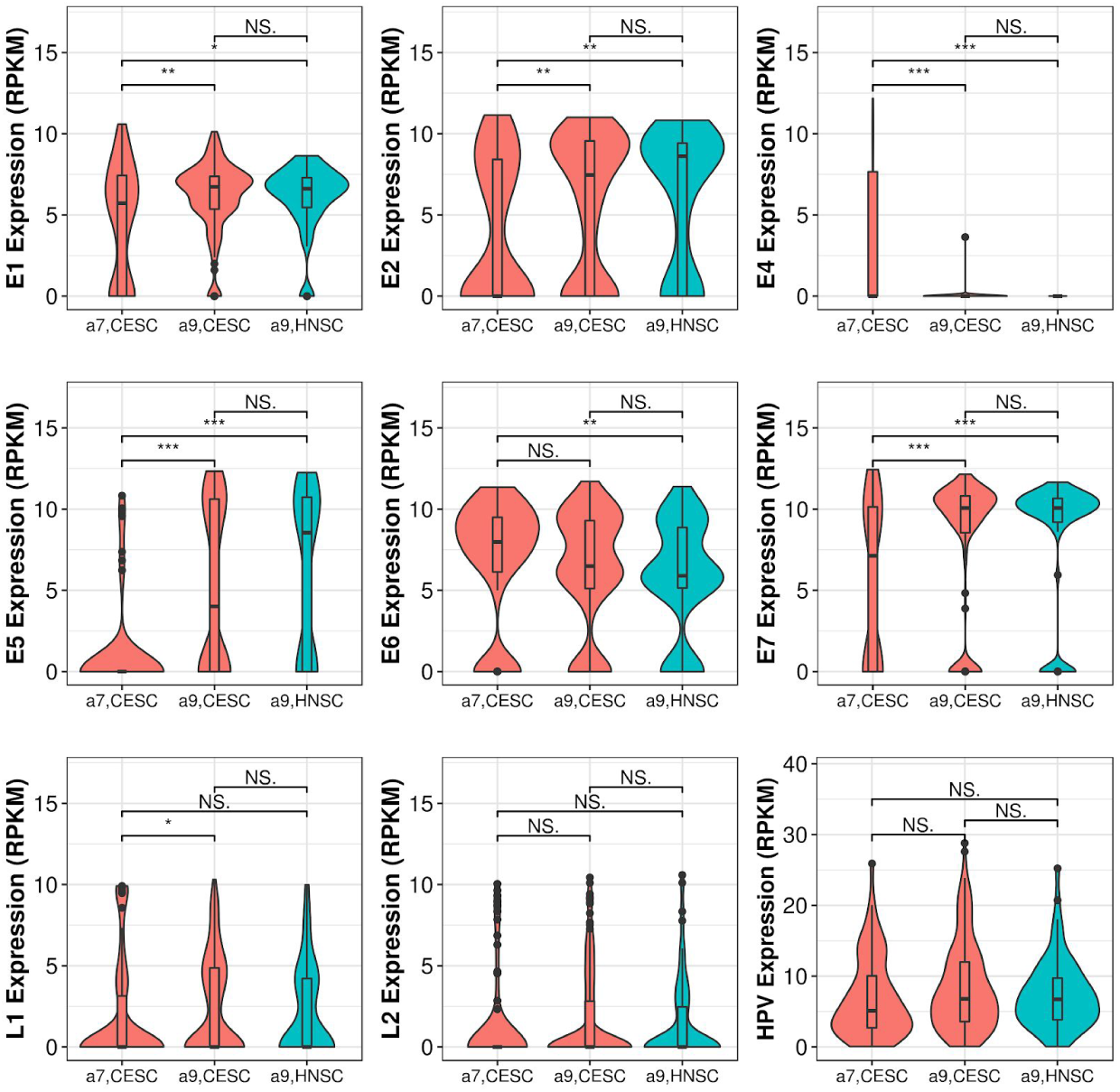
HPV gene expression varies across genotype but not cancer type. Box-and-violin plots of RPKM normalized HPV gene expression (panels 1-8) and total HPV expression in the tumor (panel 9, bottom right) between HPV a7 and a9 clades, across the TCGA cervical and head and neck cancer patients. Significant difference in gene expression between cancer types and HPV clade determined by Wilcoxon rank sum test.

## Discussion

Using multiple publically available independent datasets across three distinct oncovirally-associated cancer types, we have unraveled key correlates of HPV and HBV genotype and co-infection in oncogenic viral and host expression signaling, patient survival, tumor mutational signatures, and immune response. Apart from demonstrating that large-scale, high-fidelity *in situ* oncoviral profiling is feasible, we build upon previous studies (*24, 27*–*30*) demonstrating associations with cancer risk by showing that genotype and co-infection modalities can also affect facets of tumor evolution.

Revisiting known associations of HBV genotype C with increased risk of HCC (*30, 31*), we find that HBV genotype is also a significant predictor of overall HCC patient survival in the LIHC, with HBV genotype C driving poor survival. Remarkably, the overall effect of genotype in predicting survival is much stronger than that of tumor vascular invasion or overall tumor mutational burden, and comparable to tumor staging itself. A possible mechanistic hint for the relative lethality of genotype C lies in the significantly lowered burden of immunogenic mutations compared to genotype B. This suggests that HBV genotype B is potentially more likely to induce a host anti-tumoral immune response while HBV genotype C may evade detection and control. Additionally, this may be exacerbated by higher viral load in surrounding normal liver tissue in HBV genotype C infected patients compared with HBV genotype B infected patients (*2, 6, 32*) due to lower rates of seroclearance, compounding the effect of the more muted immunogenic tumor peptidome in HBV genotype C with an increase in viral antigen targets in the surrounding tissue. Furthermore, while no etiologically linked mutational signatures were found differentially enriched across HBV genotypes, the unique enrichment of SBS17 in HBV genotype B associated HCC has previously been correlated with increased immunogenic tumor neoantigen load (*33, 34*). However, as we found tumor expression signatures for immune markers do not correlate with HBV genotype (**Supplemental Figure 4**), this effect may play a relatively minor role in determining tumor immune infiltration. Taken together, a broader picture of preferential immune surveillance and activation in HBV genotype B over HBV genotype C tumors starts to emerge. A key caveat to these conclusions is that about 90% of patients among our HCC liver cohorts are of Asian descent, meaning only HBV genotypes B and C predominated. This necessitates a broader analysis with a more diverse patient group to confirm the trends we observe across other HBV genotypes.

Cervical tumors infected with HPV16, and more broadly strains from the a9 clade, were found enriched in tumor growth pathways over those with HPV18 (HPV a7), strongly suggesting these genotypes may be related to more aggressive tumors (*24*). The likely upregulation of *APOBEC3* in HPV16 infected tumors furthers the hypothesis of a more mutagenic tumor phenotype (*23*), although we did not find higher TMB among HPV16 patients relative to other cervical tumors. Relative enrichment of tumor hypermutation (SBS9) and defective DNA damage repair (SBS3) signatures in HPV a9 infected tumors provides further suggestive evidence for this association with HPV a9 induced tumors. Unlike HBV in HCC, however, HPV genotype on its own was not found to be a significant survival correlate. Instead, a combination of viral load, viral co-infection status, and HPV genotype significantly improved survival prediction over tumor clinical stage in cervical patients, illustrating the potential value of a more complex viral signature compared to simple infection presence or consensus genotype. Strikingly, though most cancers result from chronic infection by a single high-risk genotype of HPV (*1, 8*), we see that nearly a third of the HPV+ cervical cohort has at least two separate genotypes robustly expressed, producing an additive effect on putative tumor immunogenicity by enlarging the tumor associated peptidome of patients with multiple HPV genotypes, as seen with HPV18/HPV45 co-infected patients. Tracking tumor viral co-infections thus might identify patients who would benefit from immunotherapy or similar treatment, as has been previously investigated with intra-genotype immune-response stratification of HPV16 tumors (*35*). Whether any of these novel associations with co-infection are enhanced or diminished by inter-clade viral infections or are duplicated in other HPV associated cancers such as head and neck requires further investigation.

We attempted to address how these fine-grained properties of HPV vary across two very different tumor types by screening the HNSC TCGA cohort for HPV presence and genotype, but finding only 14% (73 patients) HPV+, of which all but one belonged to the a9 clade. Enrichments in mutational signatures linked to DNA damage repair and hypermutation (SBS3,9) found in HPV a9 cervical cancers were likewise enriched for HPV a9 head and neck cancers, while SBS40 present in the HPV a7 cervical cancer was similarly absent. This indicates that these phenotypes may be related to HPV infection, and specifically to infections of the a9 clade, even across cell-types. Furthermore, we noted that while overall HPV expression was not significantly different across cervical and head and neck cancers, significant differences in expression were observed between viral clades at the viral gene level, especially in oncovirally linked genes *E7* and *E2*. As cervical tumors have typically higher reported viral integration rates than head and neck tumors (*14, 36*–*38*), these differences may partially be explained by rates of HPV integration, which has been shown to decrease *E2* and *E5* expression. However, as gene expression profiles across the same HPV clade but different cancer types do not significantly vary, this further suggests HPV genotype plays an outsized role in defining virally linked molecular signatures across disparate cell-types.

In conclusion, we present a detailed view of key cancer types through the lens of their oncoviral cofactors by applying a scalable, highly accurate method of DNA viral gene expression, genotyping, and co-infection typing on *in situ* tumor RNA-seq data. By easily applying our method to large publically available datasets, we demonstrate that our initial analyses are sufficiently powered to reveal novel genotype, co-infection, and viral gene expression specific signals, which we hope can be built upon to help inform the next generation of clinical guidelines in assessing and treating not only HPV and HBV related cancers but other DNA virus (e.g. Epstein-Barr and Burkitt’s lymphoma) related tumors.

## Materials and Methods

### RNA-Seq analysis

RNA-Seq, MAF files, and associated clinical data for the TCGA patients were retrieved using GDACfirehose (*firehose_get*, v0.4.13), and RNA-Seq fastq and clinical data files from GEO studies GSE94660 and GSE65485 were obtained using SRA toolkit (v2.8.0). Fastq sequences were aligned to Hg38 with STAR (v2.5.1b) in two-pass mode (*39*), and gene counts for Gencode v23 (www.gencodegenes.org) gene annotated counts were generated using *featureCounts*. Mapped reads were used to allelotype (MHC class-I loci) each patient (*40*) and estimate the putative TIL burden per patient by profiling TCR and BCR sequences with MiXCR (*41*), normalizing by patient library size.

### Viral genotyping with *ViralMine*

HBV and HPV genotype calls were produced using *ViralMine* (https://github.com/LosicLab/ViralMine). Briefly, raw RNA sequencing reads that did not map to the GRCh38 reference genome were assembled into contigs using Trinity (using *–no_run_chrysalis –no_run_butterfly* flags, which only invokes the Inchworm step) to perform greedy kmer-25 contig assembly (*42*). Resulting contigs at least 100 bp long and supported by at least 20 reads were retained and CD-Hit was used to consolidate highly similar contig sequences (e-values < 1e-10) (*43, 44*). Contigs were then aligned with BLAST (*-max_target_seqs 5, -evalue 1e-6*) against HBVdb (*45*) reference genomes (genotypes A-F, and G) for HCC patients, and against RefSeq HPV reference genomes (genotypes HPV16, 18, 31, 33, 35, 45, 51, 52, 56, 58, 59, 68, 73, and 82) for cervical and head and neck cancers, effectively identifying the viral positive tumors. Viral contigs were realigned to respective viral reference genomes using a sliding window (300bp) BLAST and alignment bitscores for each window tallied by contig. Patient viral genotypes were assigned by selecting the reference genotype with the highest total combined bitscores across viral matching contigs. Patients with more than one genotype receiving greater than 10% of the bitscore total were called “co-infected”, and considered infected with all genotypes reaching this threshold.

### Differential expression and gene set enrichment analysis

Human genes from STAR *featureCounts* (see RNA-Seq analysis above) were filtered to genes with 15 or more reads across all samples in a given cohort, followed by library size normalization and expression model fitting using a negative binomial distribution with DESeq2 (*46*). Differentially expressed genes (DEGs) were identified using an absolute log-fold change < 2 and a FDR < 0.01. Design and contrast matrices were constructed based on sample comparison groups, and gene set enrichment of resulting DEGs was performed using ‘camera’ from limma analysis package for R (*47*). The top 15 enriched pathways for each analysis, by p-value, were retained used for barcode plotting (**Figures 2,3; Supplemental Table 1**).

### Integration site analysis

Viral reference genomes (HBV genotypes A, B, and C, and all tested HPV genotypes) from HBVdb (*45*) and RefSeq were included in a modified hg38 reference genome GTF as new chromosomes. Patient RNA sequences were then aligned to the respective references with STAR (v2.5.1b) in two-pass mode, and fusion junctions generated (flags *--chimSegmentMin 20 --chimJunctionOverhangMin 20 --chimScoreSeparation 6*). Large gapped chimeric alignments were filtered out by CIGAR string. For the HCC cohorts, insertions were called with a minimum of 3 supporting reads using STARchip (https://github.com/LosicLab/starchip) (*48*) (STARchip detection parameters *splitReads 3, uniqueReads 2, spancutoff 2*, and *consensus TRUE*). Due to a large perceived number of false negative integration calls in the LIHC cohort, the same fusion parameters were used without the consensus flag (*consensus FALSE*). For patients from the CESC TCGA, insertions were called with a minimum of 2 supporting reads with STARchip. Viral integration sites were compiled by limiting the resulting fusion calls to those involving only viral reference chromosomes and identifying the nearest human gene partner and sequence position of the fusion, by patient.

### Survival modeling

Cox proportional hazard models were constructed using the ‘cph’ function from the rms package (*49*), with model feature coefficients calculated using glmnet (alpha = 0.5). Samples with missing survival times, viral genotypes, and tumor clinical stage were removed, while other missing feature data was imputed. Relative predictive power of survival models against others in the same cohort was assessed using the ‘pec’ R package, computing Integrated Brier scores with .632+ bootstrap resampling across 200 iterations (*B*=200). The weights of the Brier scores correspond to the probability of not being censored, and the Kaplan-Meier estimator was used for censoring times in each resampling. Model prediction error curves were compared to the reference and other model curves for Brier scores at times comprising approximately 80% of total patient events. Test significance is indicated in the plot tables (**Figures 2D, 3C**).

### Tumor mutational signature profiling

Mutations and RNA-Seq data for tumors from each TCGA cohort were stratified by viral genotype and *de novo* mutational signatures generated using sigProfiler (https://github.com/AlexandrovLab) (*25*). Comparisons of the single-base substitution (SBS) *de novo* signatures with established SBS signatures from COSMIC (v3, https://cancer.sanger.ac.uk/cosmic/signatures/SBS/) identified group level enrichments for each cancer type. Percent SBS enrichments in each established signature were reported for each viral subgroup using the average enrichment across patient tumors in each group. Groups were considered enriched in a given SBS signature given one *de novo* profile contained a >0.5% match with the established SBS signature and the signature was present at an average of >1% across patients in the group. Linked etiologies for each SBS signature found in **Figure 4** are sourced from COSMIC.

### Tumor neoantigen prediction

To predict patient neoantigen burden and immunogenicity, we used Topiary (*50*) (https://github.com/openvax/topiary) to call mutation-derived cancer T-cell neoepitopes from somatic variants (from MAF files), tumor RNA-seq data, and patient MHC class-I HLA type. This tool matches mutations with gene annotations, filters out non-protein coding changes, and finally creates a window around amino acid changes, which is then fed into netMHCpan for each patient HLA allele across tiles of 9-12 amino-acids in length. Given that HLA-I processes neoantigens by degradation to non-conformational 8-11 amino-acid residues, we excluded neoepitopes with mutations obscured to T-cells within HLA-I binding pockets. In the case of frameshift mutations, in principle this window starts from the mutation minus the length of the peptide up to the first stop codon. Predictions were filtered by a binding affinity threshold of ic50 < 5000nM.

### Viral gene expression analysis

HPV contigs extracted from patient tumor RNA-Seq were aligned to HPV reference genes for genotypes HPV16 and HPV18 from RefSeq (builds NC_001526.4, NC_001357.1), as part of the ViralMine pipeline (https://github.com/LosicLab/ViralMine). These genotypes were used for gene expression quantification for all HPV clade a7 and a9 infected tumors, as validated HPV reference gene sequences for rarer genotypes were not readily available. Read counts for each viral gene were assigned using the read support for aligned viral contig. HPV genes with no patient contig alignment were considered not expressed and given read counts of 0. HPV gene expression was RPKM normalized using patient tumor RNA-Seq library size.

### Statistical Analysis

The effect of read depth threshold tumor on viral genotyping with ViralMine was determined by subsampling RNA-Seq reads from the cervical and HCC patients at 95, 90, 75, 50, 25, 10, 5, 1% of total reads (**Supplemental Figure 9**). While viral genotype true positives were consistently confidently called at low subsampling rates, subsampling below 25% of total reads saw an increase in the number of false positives and false negative calls, in both patient groups. Therefore, any samples with total coverage less than 25% of the average coverage across each cohort would be removed from analysis. As this threshold was low and sample coverage relatively uniform across patients, no samples from any cohort were removed by this filter. An additional head and neck cancer immunotherapy treated cohort from *Miao et al.* (*51*) was mined to understand the effects of anti-CTLA-4/PD-1 therapy on viral contig recovery (**Supplemental Figure 10**). Though there were only 4 HPV positive samples in the immunotherapy cohort, the number of viral reads recovered in these patients was significantly fewer than patients in TCGA HNSC cohort, signaling immunotherapy may significantly reduce viral RNA in the tumor and make viral detection and genotyping more difficult.

The co-infection threshold was determined by comparing the number of samples with co-infections given a varying bitscore fractional limit (1-50% of bitscore total) for calling a given HPV or HBV genotype as detected (**Supplemental Figure 11**). An inflection point in the relationship between co-infection total and bitscore alignment fraction cutoff was observed at 0.1 (or 10%), at which point further cutoff reduction saw an exponential increase in the number of samples with co-infections. As such, 0.1 was selected as the limit for co-infection detection.

Survival model prediction error curve Brier scores were compared to reference curves using a one-sided Kolmogorov-Smirnov test (ks-test), since scores are not normally distributed and we are interested in determining the relative significance of one model over the reference. We used a likelihood ratio test to determine the relative significance of model terms between nested models.

For patient-level mutational signature (SBS) composition profiling, *k*-means clustering was used to stratify patients by total viral expression and TMB (**Figure 4b, Supplemental Figure 6**). The number of clusters for each viral subgroup was set at 1/4th the total group size, with a lower threshold of 25 patients, in which case the entire group was represented. Single patients were sampled without replacement from each cluster to form a representative set for each viral subtype. Associations between total viral expression, TMB and signature enrichment were determined by linear modeling of SBS signatures by stratified features, with associated coefficient p-values reported.

Comparisons between tumor neoantigen cumulative distributions were made using a one-sided ks-test. Comparison of the ratio of immunogenic mutation burden between viral genotype groups was done using Wilcoxon test. One sample from each of the HPV18 and HPV45 patient groups was removed before from comparison of immunogenic mutational burden with the co-infected HPV18/45 group, as presented in **Figure 5**. Samples were removed as major outliers violating the general distribution of ratios. Finally, normalized HPV gene expression was compared between genotype and disease type groups using the Wilcoxon test, chosen due to the non-Gaussian distribution of expression levels. Significance was considered at p < 0.05 for all tests.

## Data and Code Availability

RNA-Sequencing data is available through GDAC and GEO as referenced in the methods. ViralMine and associated viral reference databases are available for download (https://github.com/LosicLab/ViralMine). Additional analysis code is available upon reasonable request to the corresponding author.

## Acknowledgements

This work was supported in part through the computational resources and staff expertise provided by Scientific Computing at the Icahn School of Medicine at Mount Sinai.

