## Supplemental Figures for "Landscape of oncoviral genotype and co-infection via Human papilloma and Hepatitis B viral tumor in-situ profiling"

*[1]Departments of Genetics and Genomic Sciences, [2]Division of Liver Diseases, Division of Hematology/Oncology, Department of Medicine, Graduate School of Biomedical Sciences, Tisch Cancer Institute, Diabetes, Obesity, and Metabolism Institute, [3]Department of Otolaryngology Head and Neck Surgery, [4]Division of Hematology Oncology, Department of Medicine, Icahn School of Medicine at Mount Sinai, New York, NY, 10029*
*[5]INSERM, U1052, Cancer Research Center of Lyon (CRCL), Lyon, France, 69008*

### Supplementary Materials:

1. **Supplemental Fig. 1**: Sensitivity and specificity of ViralMine in validation cohorts.
2. **Supplemental Fig. 2**: Cervical cancer histology does not correlate with HPV infection
3. **Supplemental Fig. 3:** Co-infection rates of HPV and HBV in related cancers of the TCGA
4. **Supplemental Fig. 4:** Viral genotype and viral load not associated with other clinical covariates or immune measures
5. **Supplemental Fig. 5**: HCC HBV co-infection v. single infection GSEA results
6. **Supplemental Fig. 6:** HBV integration sites by HCC cohort
7. **Supplemental Fig. 7:** Recurrent integration sites of HPV in CESC TCGA patients
8. **Supplemental Fig. 8:** SBS mutational profiles of patients with viral infected tumors organized by TMB
9. **Supplemental Fig. 9**: Accuracy of ViralMine over subsampling of RNA-Seq reads
10. **Supplemental Fig. 10**: HPV Reads for head and neck tumors treated with immunotherapy and not treated (TCGA)
11. **Supplemental Fig. 11**: Single-virus co-infections called given variable threshold
12. **Supplemental Table 1**: Top CAMERA gene set enrichment results for LIHC and CESC comparisons
13. **Supplemental Table 2**: Mutational signature average enrichments by HPV and HBV genotype
14. **Supplemental Table 3**: HPV associated TCGA head and neck tumors by anatomical site

### Supplemental Figures

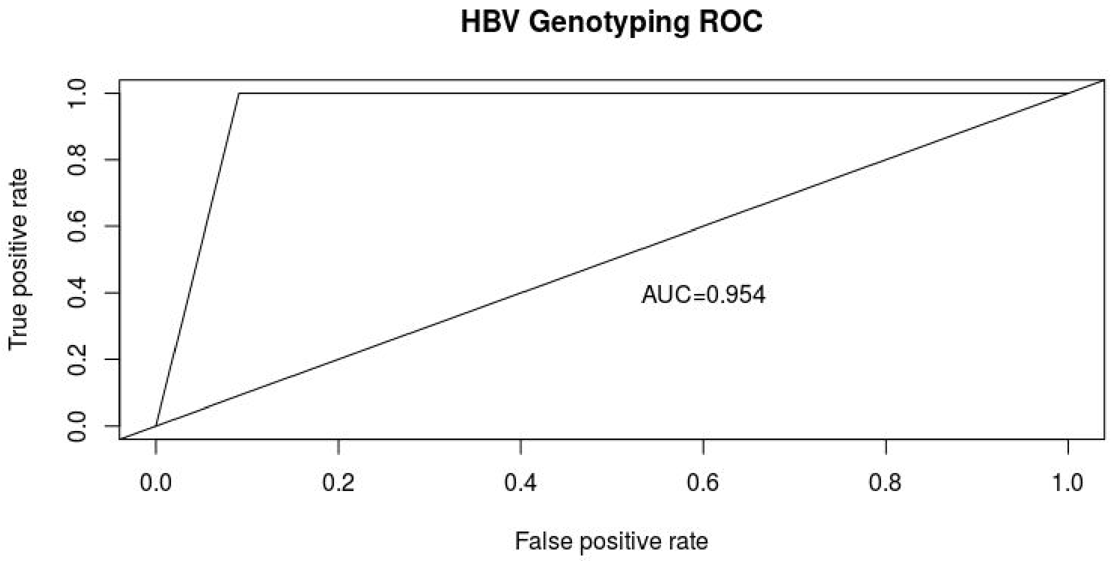


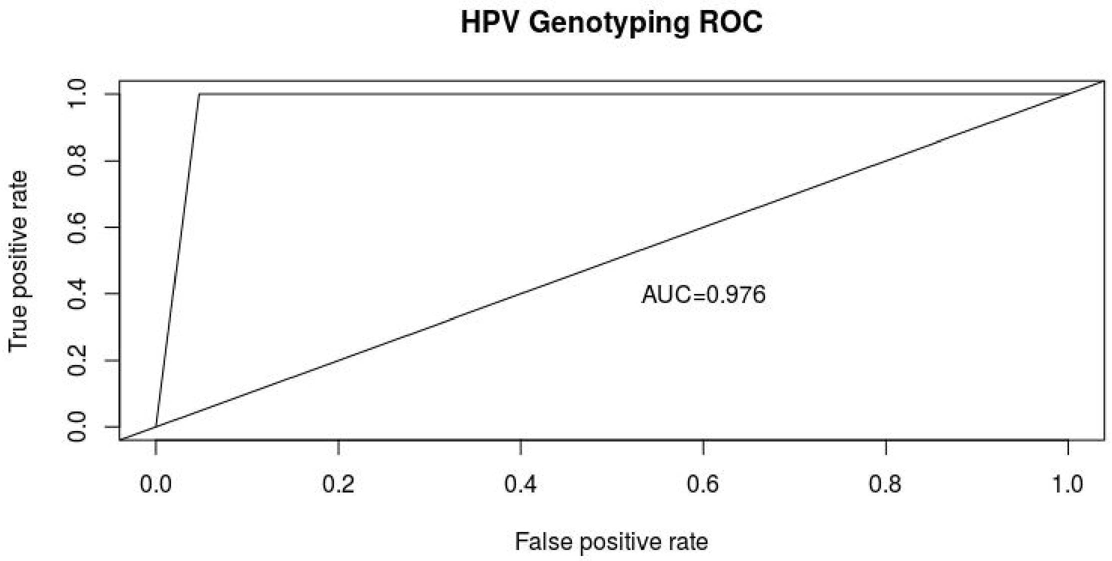


**Supplemental Fig. 1: Sensitivity and specificity of ViralMine in validation cohorts.** ROC curves for HBV genotyping in the GSE65485 HCC cohort (top) and the core set of HPV+ patients in the TCGA cervical cancers (bottom). True positives were called if viral genotype called by our method matched the genotype reported by the respective study, while false positives were called if ViralMine reported a viral genotype for a sample reported previously as virally negative.

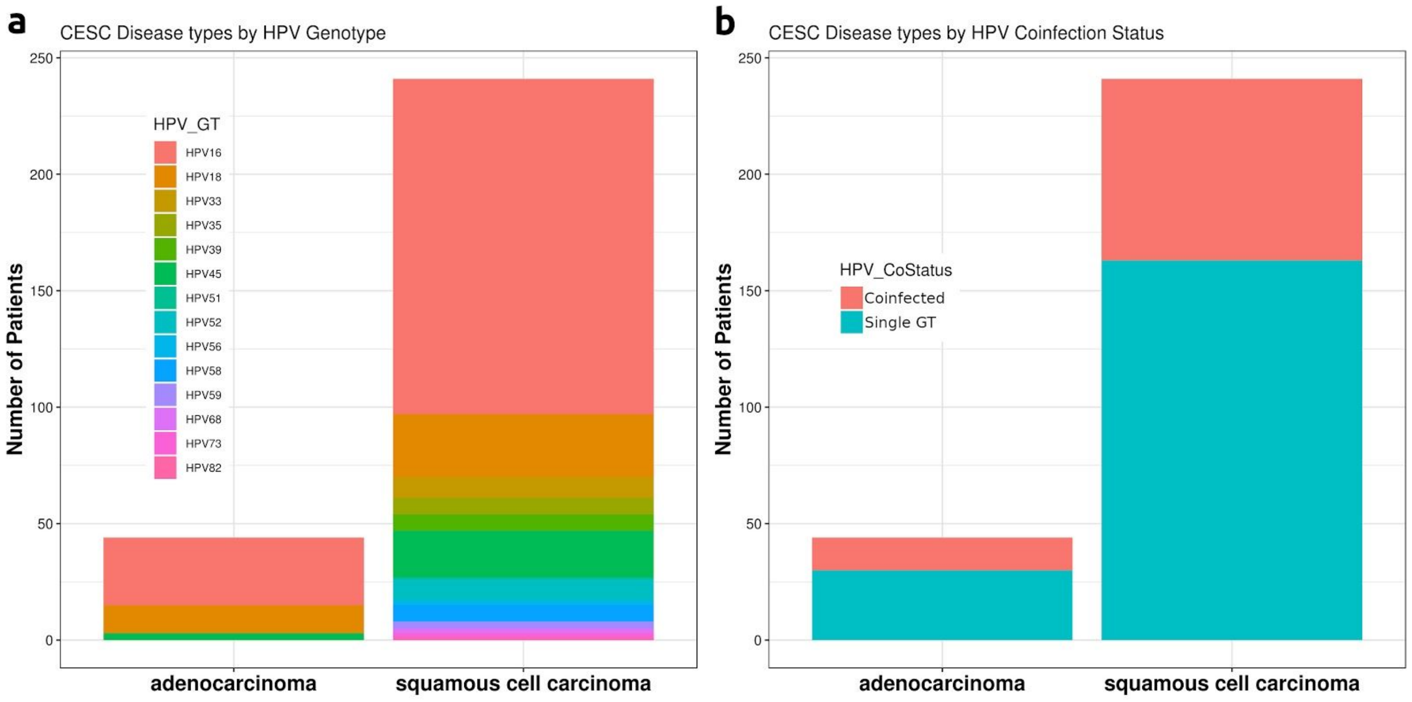


**Supplemental Fig. 2: Cervical cancer histology does not correlate with HPV infection.** HPV+ TCGA cervical cancer patients (CESC) divided by broad disease histology show no significant association with (A) HPV genotype, or (B) co-infection of multiple HPV genotypes, controlling for group sample size.

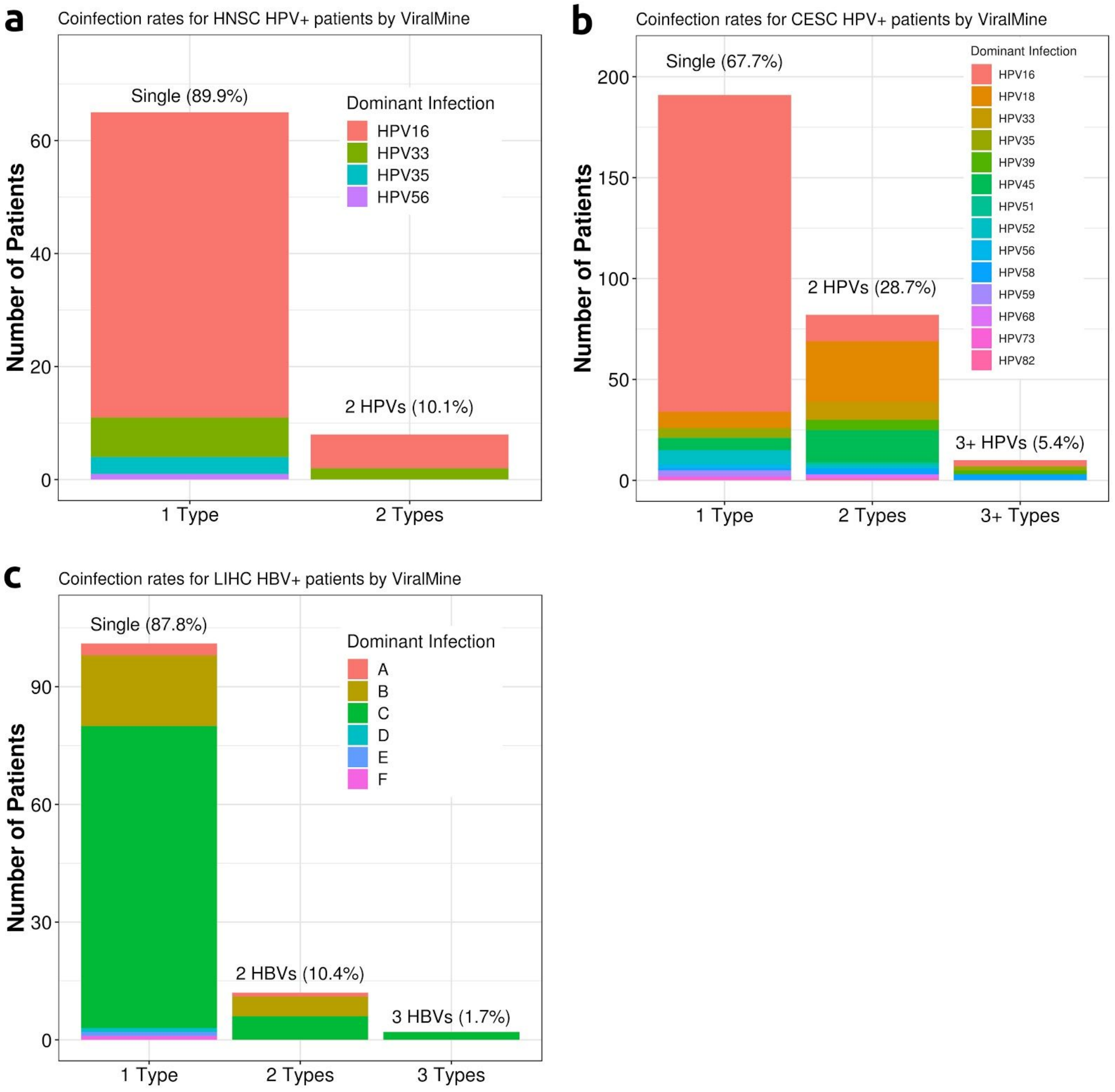


**Supplemental Fig. 3: Co-infection rates of HPV and HBV in related cancers of the TCGA.** Patients are divided by number of viral co-infections of HPV or HBV present and colored by dominant detected genotype of infection for (A) cervical cancer, (B) head and neck cancers, and (C) liver cancer.

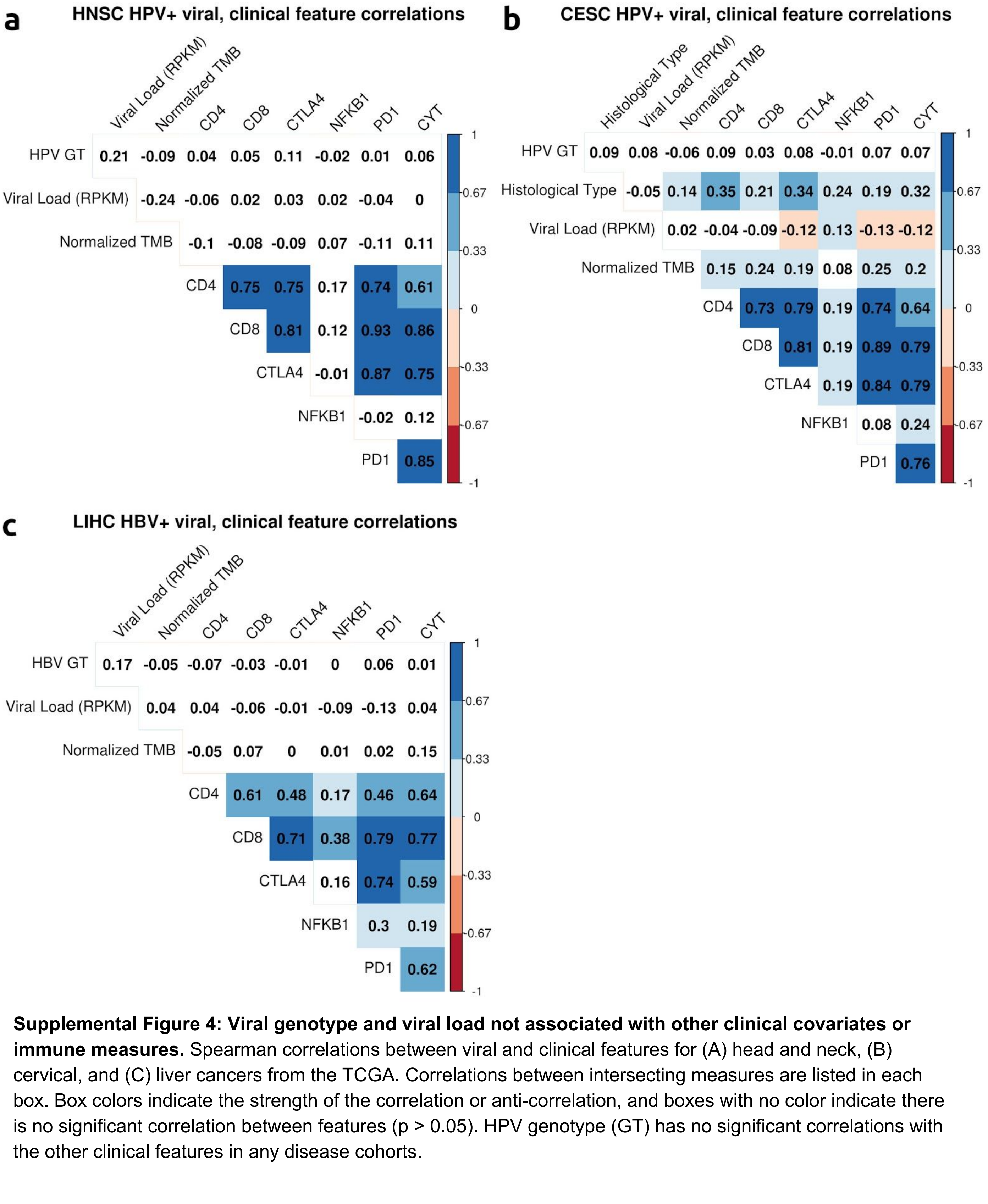


**Supplemental Figure 4: Viral genotype and viral load not associated with other clinical covariates or immune measures.** Spearman correlations between viral and clinical features for (A) head and neck, (B) cervical, and (C) liver cancers from the TCGA. Correlations between intersecting measures are listed in each box. Box colors indicate the strength of the correlation or anti-correlation, and boxes with no color indicate there is no significant correlation between features ($p > 0.05$). HPV genotype (GT) has no significant correlations with the other clinical features in any disease cohorts.

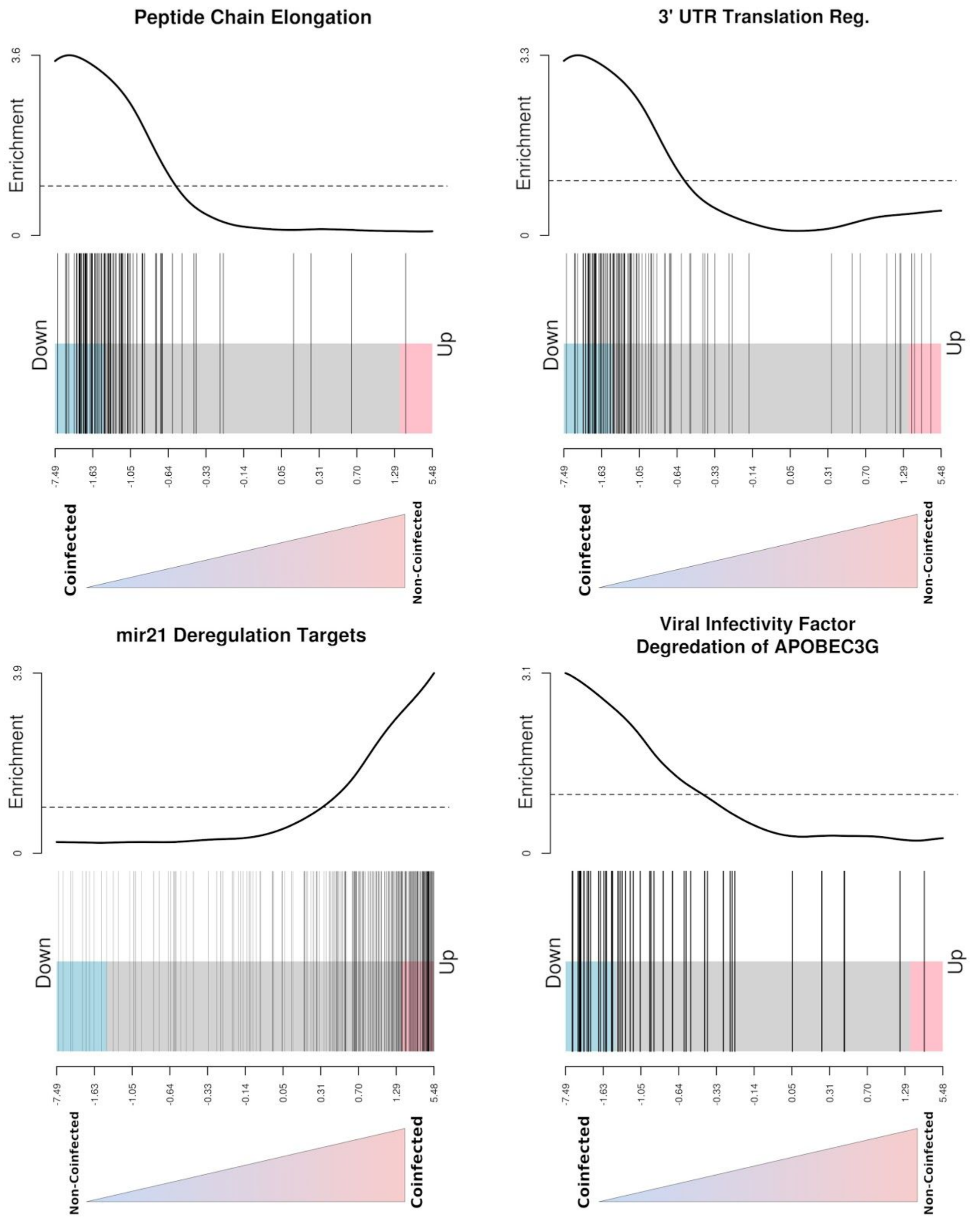


**Supplemental Fig. 5: HCC HBV co-infection v. single infection GSEA results.** Barcode plots indicating gene up- and downregulation between HBV co-infected and single-genotype infected liver tumors. Cell regulatory pathways in peptide chain elongation and 3' UTR translation are down expressed with co-infection, as are mir21 deregulation targets with respect to single-infected like tumors. Additionally APOBEC degradation appears to down expressed in co-infected patients. Bars along the bottom of the plot indicate individual pathway gene enrichment scores.

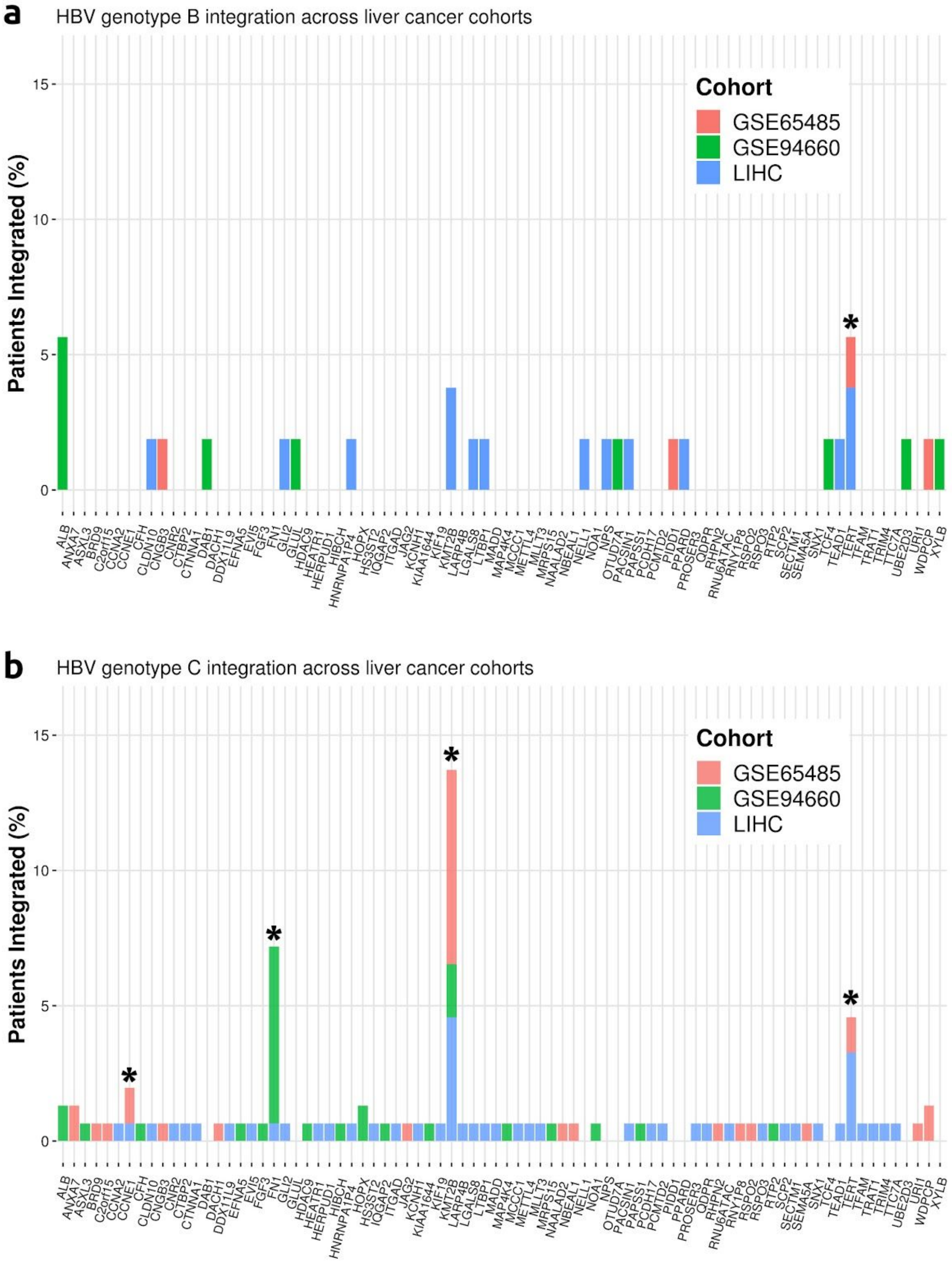


**Supplemental Fig. 6: HBV integration sites by HCC cohort.** Integration sites found by StarChip Fusion for (A) HBV B associated and, (B) HBV C associated HCC. Genes recurrently integrated across HCC cohorts are indicated with an asterisk. Sites in pseudo-genes have been removed for clarity.

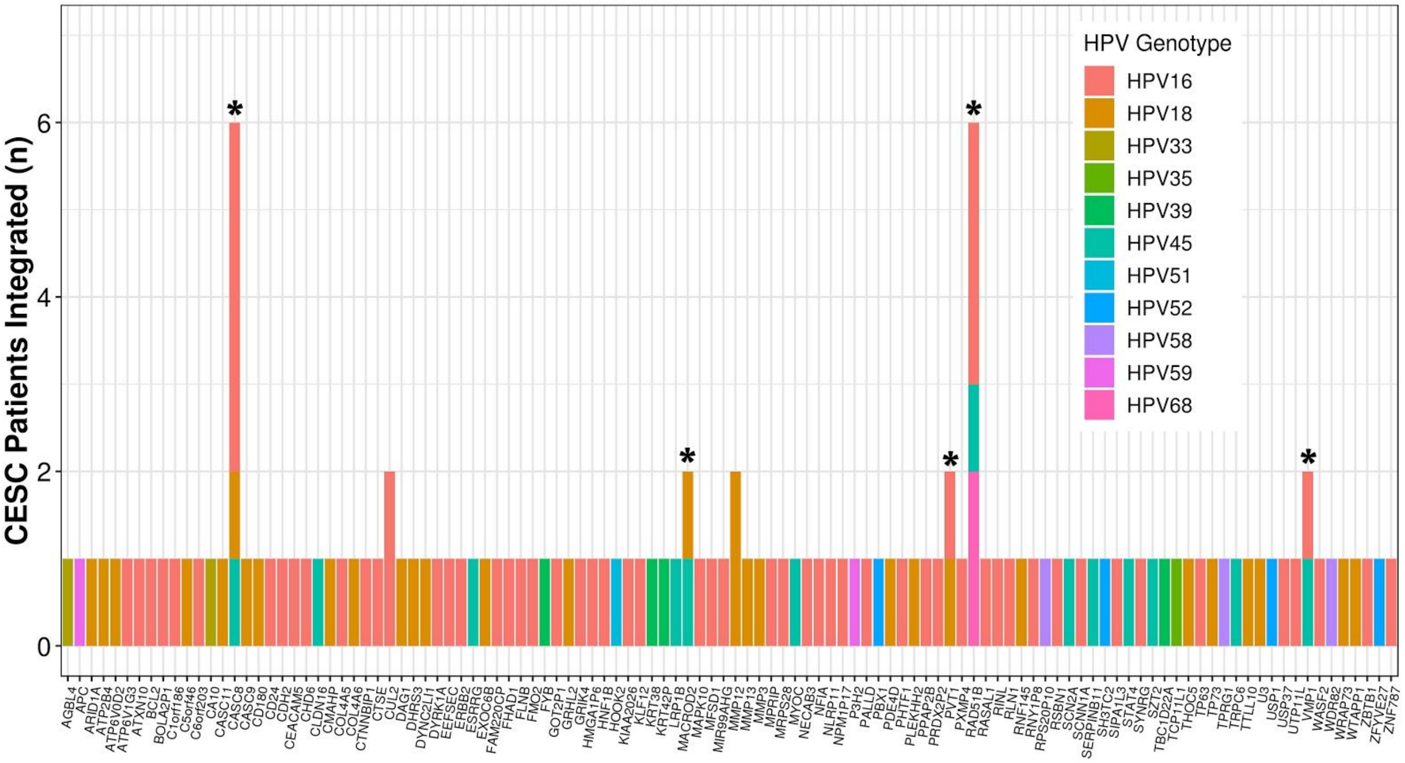


**Supplemental Fig. 7: Recurrent integration sites of HPV in CESC TCGA patients.** HPV Integration sites found by StarChip Fusion across cervical tumors in the TCGA cohort. Genes recurrently integrated across HCC cohorts are indicated with an asterisk. Sites in pseudo-genes have been removed for clarity.

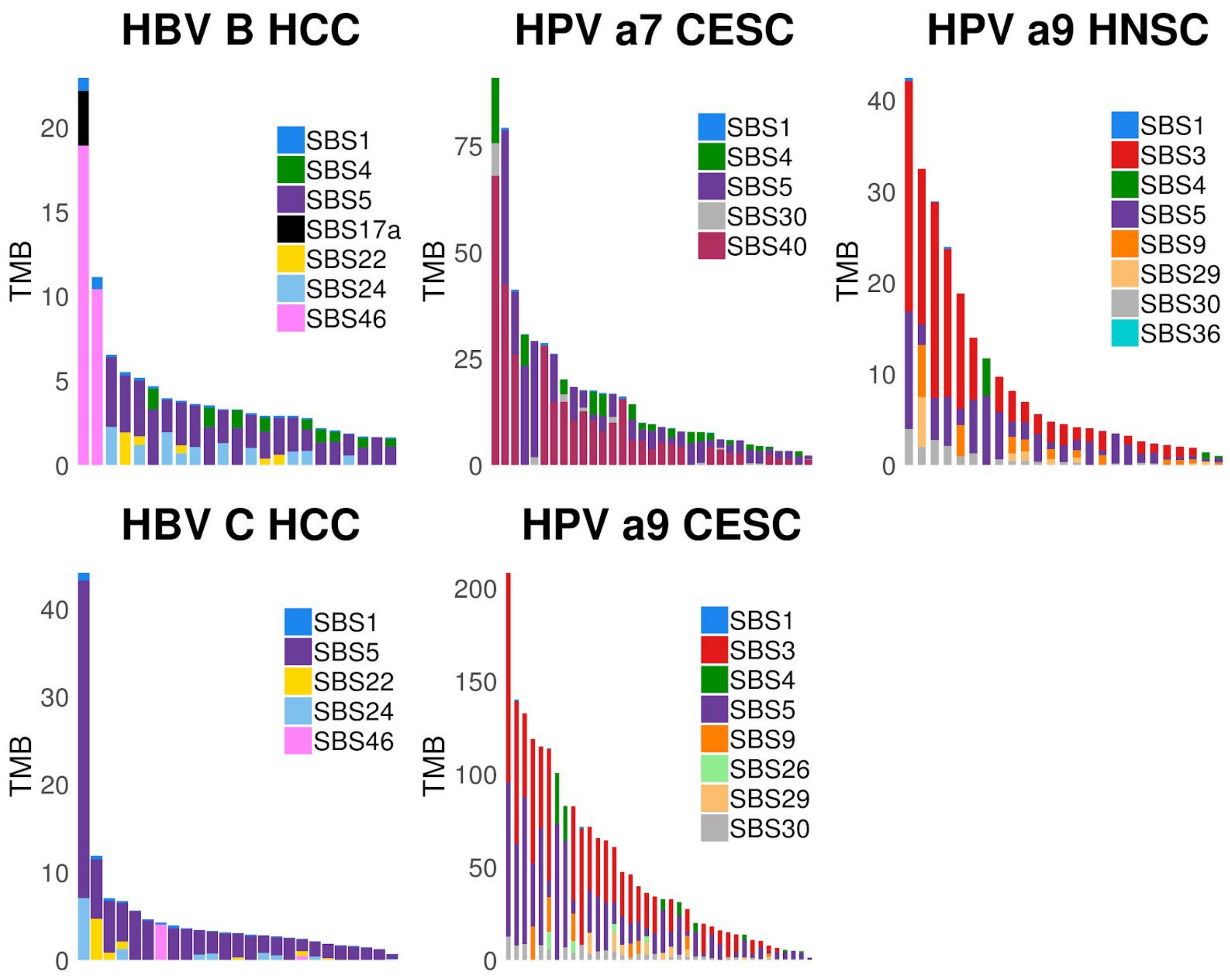


**Supplemental Fig. 8: SBS mutational profiles of patients with viral infected tumors organized by TMB.** Signature contributions across representative patient mutational profiles, organized by tumor mutational burden. Each bar represents one patient of the given viral genotype group, with color fill corresponding to the proportion of the patient's total mutations attributed to the indicated SBS signature.

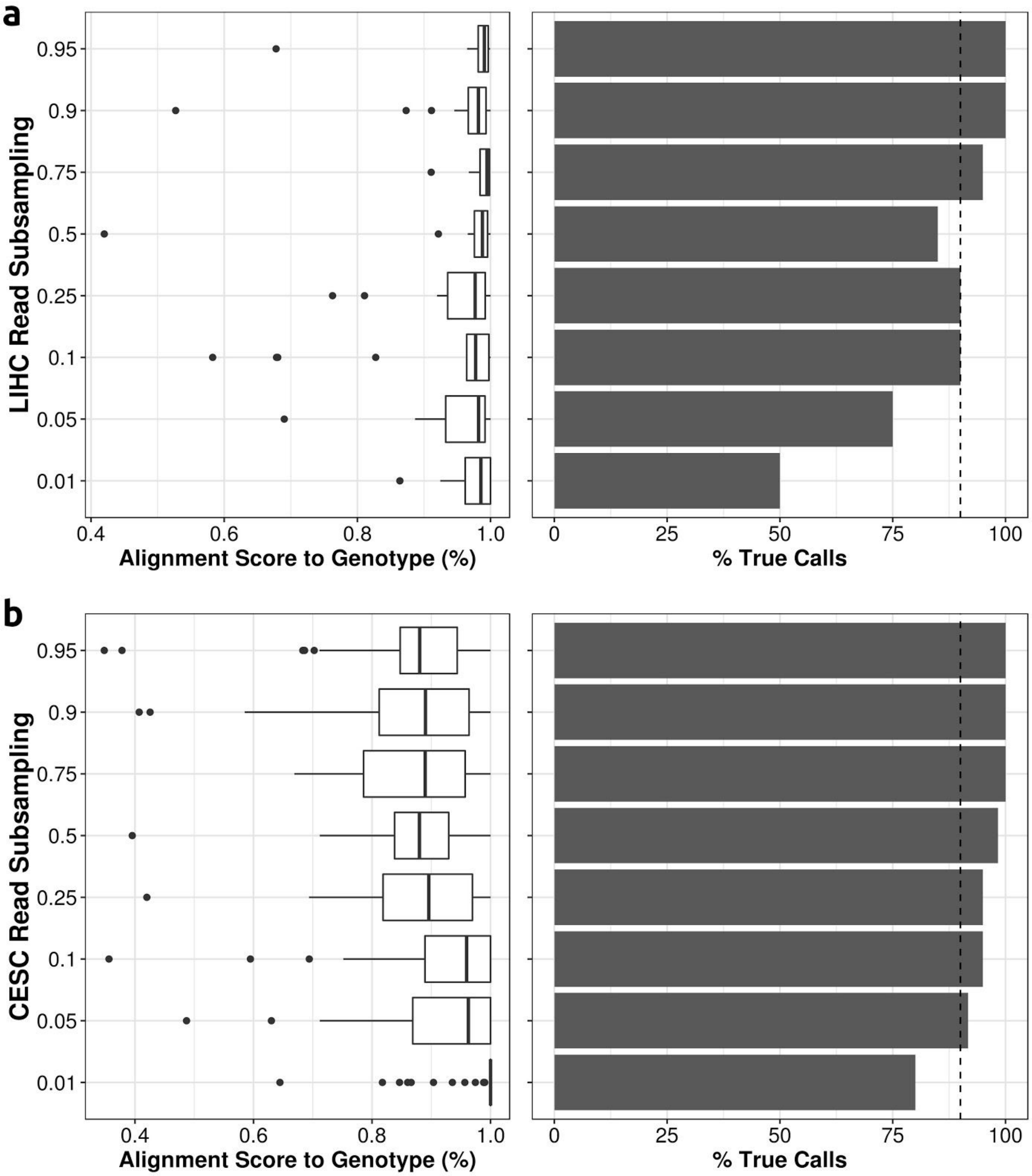


**Supplemental Fig. 9: Accuracy of ViralMine over subsampling of RNA-Seq reads.** Boxplots (left) of the percentage alignment score total captured by true positive viral genotype calls, given read subsampling rate (y-axis) across randomly sampled patients from the (A) HBV+ LIHC cohort, and (B) HPV+ CESC cohort. Rate of true HBV genotype calling at each read subsampling level presented in the adjacent barplots. Dotted vertical lines indicate 90% true positive call rate.

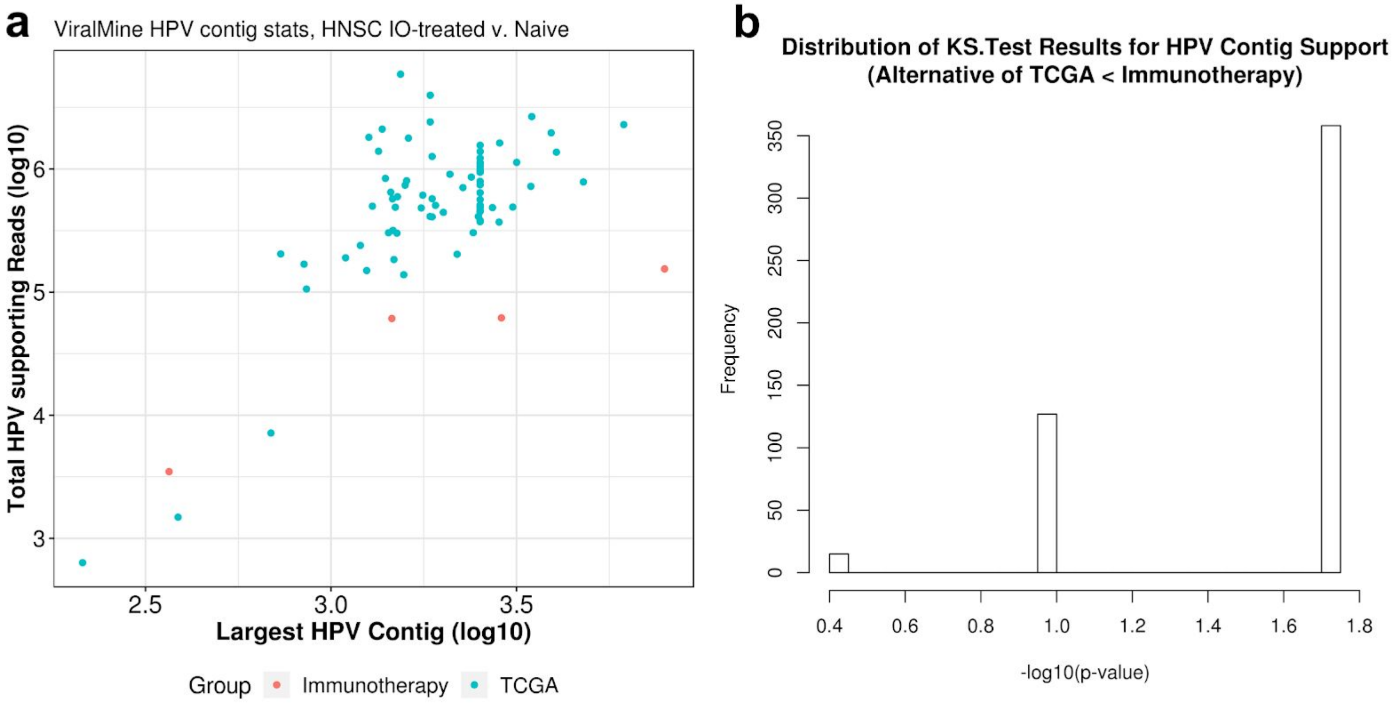


**Supplemental Fig. 10: HPV Reads for head and neck tumors treated with immunotherapy and not treated (TCGA).** (A) Largest recovered HPV contigs from ViralMine against total HPV contig supporting reads for HPV+ head and neck patients from the TCGA and patients treated with anti-CTLA-4/PD-1 (*Van Allen*, 2018). (B) Distribution of KS-Test p-values comparing distribution of total HPV supporting reads between immunotherapy treated patients and 4 randomly subsampled TCGA patients, over 500 replicates. Distribution comparisons generally indicate significantly fewer HPV reads for immunotherapy treated patients than TCGA patients.

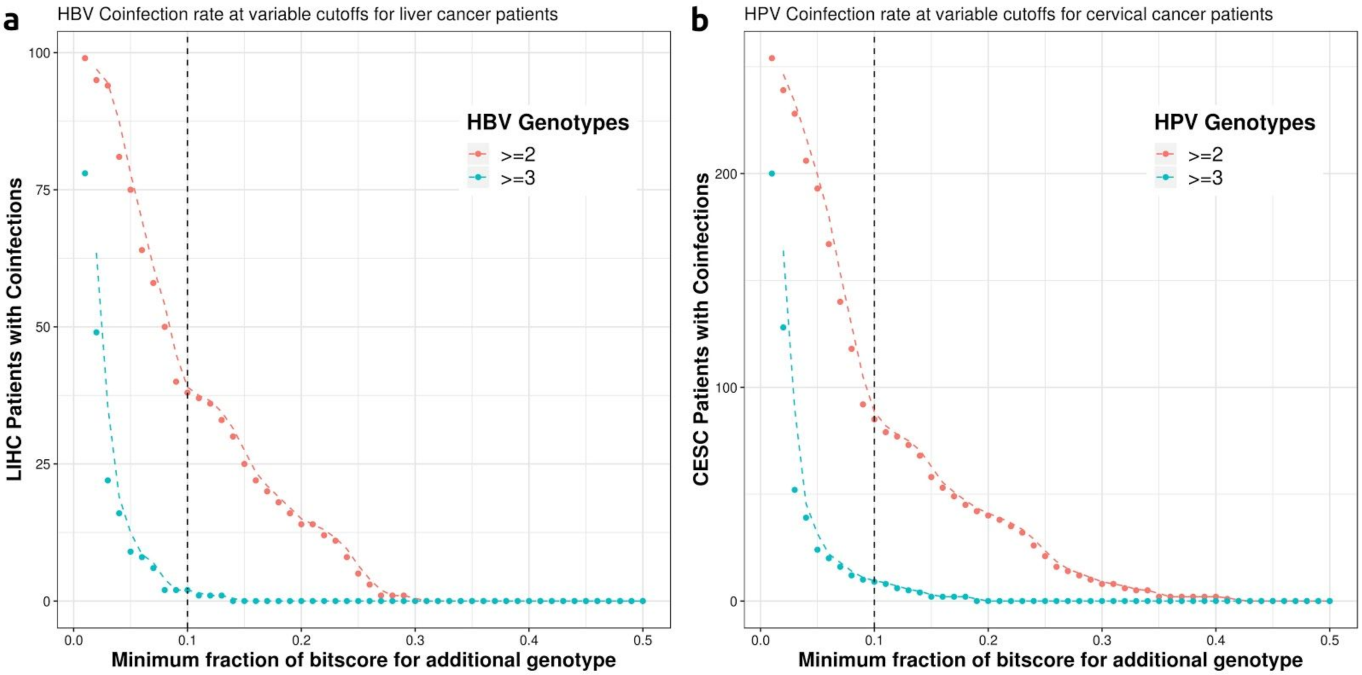


**Supplemental Fig. 11: Single-virus co-infections called given variable threshold.** Number of viral genotypes called for (A) HBV positive HCC patients from the TCGA LIHC, and (B) HPV positive cervical cancer patients from the TCGA CESC after adjusting the biscore alignment threshold used to determine single-virus co-infection. An inflection point in the number of patients with two or more genotypes detected is observed at an alignment fraction threshold of 0.1, or 10% of the bitscore total.
